# Induction of a colitogenic phenotype in Th1 cells depends on IL-23R signaling

**DOI:** 10.1101/2021.01.24.426445

**Authors:** Mathias Pawlak, David DeTomaso, Gerd Meyer zu Horste, Youjin Lee, Jackson Nyman, Danielle Dionne, Chao Wang, Antonia Wallrapp, Patrick R. Burkett, Samantha J. Riesenfeld, Ana C. Anderson, Aviv Regev, Ramnik J. Xavier, Nir Yosef, Vijay K. Kuchroo

**Author notes:** Department of Neurology, Institute of Translational Neurology, Medical Faculty, University Hospital Muenster, Germany (Gerd Meyer zu Horste). Pritzker School of Molecular Engineering & Biological Sciences Division, Genetic Medicine, University of Chicago, IL. Department of Microbiology and Immunology and Diabetes Center, University of California, San Francisco, CA, USA; Innovative Genomics Institute, University of California, Berkeley, CA, USA. Genentech, South San Francisco, CA 94080, USA. Contributed equally (Mathias Pawlak, David DeTomaso).

## Abstract

The cytokine receptor IL-23R plays a fundamental role in inflammation and autoimmunity. However, several observations have been difficult to reconcile under the assumption that only Th17 cells critically depend on IL-23 to acquire a pathogenic phenotype. Here, we report that Th1 cells differentiated *in vitro* with IL-12 + IL-21 show similar levels of IL-23R expression as in pathogenic Th17 cells. We demonstrate that IL-23R is required for Th1 cells to acquire a highly colitogenic phenotype. scRNAseq analysis of intestinal T cells enabled us to identify novel regulators induced by IL-23R-signaling in Th1 cells which differed from those expressed in Th17 cells. The perturbation of one of these regulators (CD160) in Th1 cells inhibited induction of colitis. In this process, we were able to uncouple IL-23R as a purely Th17 cell-specific factor and implicate IL-23R signaling as a pathogenic driver of Th1 cell-mediated tissue inflammation and disease.

## Introduction

The cytokine IL-23 and its receptor IL-23R play a fundamental role in inducing tissue inflammation and autoimmunity (Teng et al., 2015). It has been shown that pre-clinical models of Multiple Sclerosis, arthritis and inflammatory bowel disease (IBD) are all protected from disease in animals deficient for IL-23 signaling (Ahern et al., 2010; Cua et al., 2003; Murphy et al., 2003). The relevance to human disease is emphasized by genome-wide association studies (GWAS) that established *IL-23R* as a risk gene in multiple human autoimmune diseases including IBD (Duerr et al., 2006) and a number of anti-IL-23 inhibitors have been approved for the treatment of Psoriasis and are now being tested in other autoimmune conditions where Th17 cells have not been implicated in disease induction (Patel and Kuchroo, 2015).

IL-23R signaling is crucial for evoking a pathogenic phenotype in Th17 cells by stabilizing their function and inducing multiple factors that make Th17 cells highly pro-inflammatory (McGeachy et al., 2009). Although the Th17 cell subset is most prominently associated with IL-23R function, several observations, however, have been difficult to reconcile under the assumption that IL-23R solely operates in pathogenic Th17 cells and not other pro-inflammatory T cell subsets. Importantly, in the pre-clinical colitis models, IL-23R signaling has also been implicated in regulating FoxP3^+^ Treg function (Izcue et al., 2008; Schiering et al., 2014). However, it has been observed that *bona fide* Th1 cells also elicit colitis in pre-clinical models, particularly in the adoptive transfer colitis model (Harbour et al., 2015), but Th1 cells are not known to express IL-23R. These data are conflicting with the observation that IL-23R signaling is required for the induction of colitis. Interestingly, a recent report suggests that Th17 cells need to transdifferentiate into Th1 cells to elicit colitis in this model (Harbour et al., 2015), however, whether Th1 cells themselves, driven by IL-23R, could trigger colitis without going through a Th17 cell-state has not been addressed. Clinically, secukinumab, a monoclonal antibody targeting IL-17A has been found effective in plaque psoriasis, psoriatic arthritis and ankylosing spondylitis yet ineffective in IBD (Hueber et al., 2012). In contrast, ustekinumab, a monoclonal antibody targeting both IL-12 and IL-23, and therefore targeting differentiation of both Th1 cells and Th17 cells, has proven to be effective in treating IBD (Sandborn et al., 2012; Sands et al., 2019b). Indeed, the seminal studies by Powrie and colleagues have established that both Th1 and Th17 cells develop in the pre-clinical disease model of IBD through the adoptive transfer of naïve CD45RB^hi^ T cells (Ahern et al., 2010; Powrie et al., 1994). These observations raised important questions and provided impetus to investigate whether IL-23 may also confer pathogenicity to another T helper cell subset, in addition to quintessential Th17 cells, to thereby contribute to the induction of IBD.

To this end, we performed a cytokine screen and discovered that the cytokine combination IL-12 + IL-21 strongly induces IL-23R on Th1 cells *in vitro*. Loss of IL-23R expression on Th1 cells almost completely abrogated the ability of Th1 cells to transfer colitis. Using massively parallel single-cell RNA-sequencing (scRNAseq) of intestinal tissue-infiltrating colitogenic Th1 cells, we demonstrated a striking expression of genes that promote inflammation and linked these with multiple pathways implicated in the induction of IBD in humans as identified by GWAS analysis. Finally, we have identified a number of novel regulators induced by IL-23R-dependent signaling in Th1 cells and have shown a critical role for the IL-23R-dependent gene *Cd160* in colitis. We thus identify a pathogenic function for IL-23R signaling in Th1 cells that promote colitis, expanding the importance of this pathway beyond Th17 cells in autoimmune disease and enabling identification of novel targets linked with human IBD for further investigation.

## Results

### *In vitro* differentiation of naïve T cells with IL-12 and IL-21 induces IL-23R^+^ Th1 cells

Although it has been shown *in vivo* that IFN-γ^+^ T cells can express IL-23R, these IFN-γ-producing cells were identified as transdifferentiated Th17 cells using a fate-mapping approach (Hirota et al., 2011). In fact, chronic stimulation of Th17 cells with IL-23 was shown to increase IFN-γ production from Th17 cells and to inhibit IL-17 production (Hirota et al., 2011). However, whether there are *bona fide* Th1 cells that express IL-23R and thus may be impacted by IL-23 signaling has not been addressed. It has been long established that naïve T cells differentiated *in vitro* with IL-12 to become Th1 cells express IL-12R but are not known to express IL-23R (Szabo et al., 2000). Therefore, we sought to identify whether there are *bona fide* Th1 cells that express IL-23R and undertook a screen to identify a cytokine that in conjunction with IL-12 would efficiently induce IL-23R in Th1 cells *in vitro*. Th1 cells express IFN-γ, IL-2, lymphotoxin (LT) and the master transcription factor T-bet but do not produce IL-17A, the signature cytokine of Th17 cells (Mosmann et al., 1986; Szabo et al., 2000). Screening different cytokine conditions (Fig. S1), we discovered that culturing naïve CD4^+^ T cells with IL-12 + IL-21 for 5 days induces strong expression of IL-23R, together with IFN-γ expression, the signature cytokine of Th1 cells (Figures 1A and S1). The expression of IL-23R in Th1 cells reached similar levels as in pathogenic Th17 cells that were differentiated with the cytokine combination IL-1β + IL-6 + IL-23 (Figure 1A) (Ghoreschi et al., 2010). Differentiation of naïve T cells with IL-12 or IL-18 alone induced minimal expression of IL-23R (Figure S1). Addition of IL-23 together with IL-12 did also induce *Il23r* expression, however, the induction of IL-23R and IFN-γ expression was more pronounced with IL-12 + IL-21 (Figure S1). We also measured the expression of *Il23r*, *Ifng*, *Il17a*, *Tbx21*, *Rorc* and *Stat3* in these *in vitro* differentiated T cells and could confirm the induction of *Il23r* together with a Th1 cell transcriptional program (Figure 1B). In summary, we demonstrate that IL-12 + IL-21 strongly induces IL-23R expression in Th1 cells *in vitro*.

**Figure 1.**
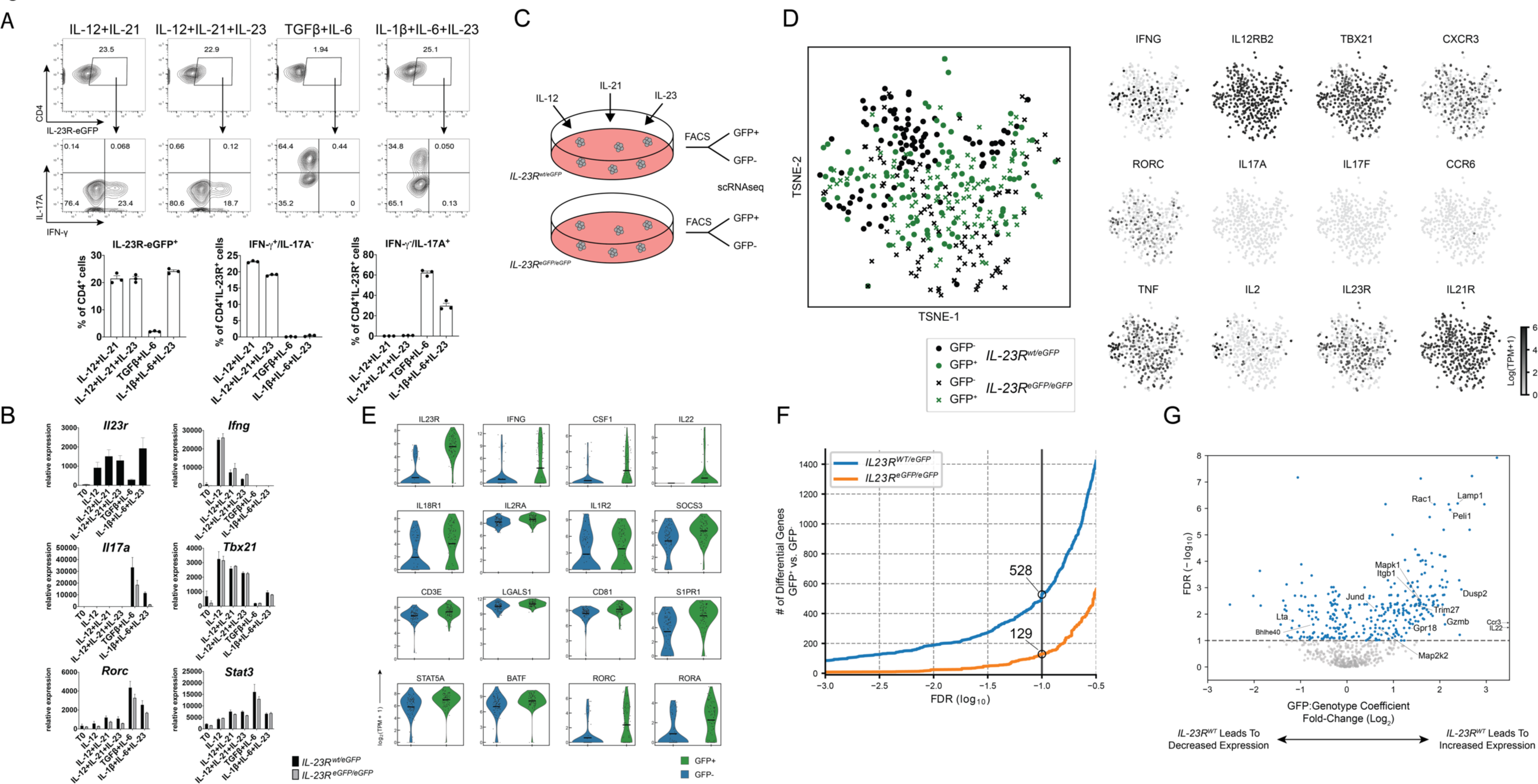
IL-12 + IL-21 strongly induce IL-23R in Th1 cells in vitro. (A) Naïve CD44^low^CD62L^hi^ CD4^+^ T cells were cultured under four different conditions *in vitro* for 5 days on anti-CD3/CD28 antibody-coated plates with the following conditions: TGF-β + IL-6 (non-pathogenic Th17 cells), IL-1β + IL-6 + IL-23 (pathogenic Th17 cells), IL-12 + IL-21 and IL-12 + IL-21 + IL-23. Flow cytometry was used to measure the extent of IL-23R expression taking advantage of an eGFP reporter allele. (B) qPCR analysis shows the expression of *Il23r* in wildtype (*Il23r^eGFP/wt^*) cells. Important signature Th1 and Th17 genes such as *Ifng*, *Il17a*, *Tbx21*, *Rorc* and *Stat3* were measured in both wildtype (*Il23r^eGFP/wt^*) and KO cells (*Il23r^eGFP/eGFP^*). (C) Schematic of sorting by FACS of IL-23R^+^ (eGFP^+^) and IL-23R^-^ (eGFP^-^) cells differentiated with IL-12 + IL-21+ IL-23 in preparation of scRNAseq (Smart-seq2). (D) t-SNE plots showing the expression of Th1 and Th17 signature genes. (E) Selected genes are shown that are expressed in eGFP^+^ and eGFP^-^ cells in wildtype (*Il23r^eGFP/wt^*) cells. Green violin plots represent eGFP^+^-sorted cells. (F) 528 genes were found to be differentially expressed between eGFP^+^ and eGFP^-^ cells from wildtype cells (*Il23r^eGFP/wt^*). In knockout cells for IL-23R (*Il23r^eGFP/eGFP^*), only 129 genes were found to be differentially expressed between eGFP^+^ and eGFP^-^ cells. (G) Volcano plot showing genes differentially expressed in an IL-23R dependent manner. An interaction coefficient was computed to identify differences between the eGFP^+^ and eGFP^-^ populations which varied according to genotype (wildtype *Il23r^eGFP/wt^* vs. knockout *Il23r^eGFP/eGFP^*). Panel (A) is representative of several experiments and mean + s.e.m are shown. In Panel (B) mean + s.e.m are shown.

### Single-cell RNA-sequencing identifies genes in Th1 cells that are expressed in an IL-23R-dependent manner

It is known that IL-23R signaling induces a set of genes in Th17 cells that makes them pathogenic, therefore, we studied the impact of IL-23R signaling on Th1 cells. We took advantage of a previously described reporter allele for *Il23r* expression and chose a two-pronged approach: (1) We differentiated naïve T cells from reporter mice that carry an eGFP reporter in their endogenous IL-23R locus (*Il23r^eGFP/wt^*) with IL-12 + IL-21 + IL-23 for 96 hours. We then isolated IL-23R^+^ (i.e. eGFP^+^) and IL-23R^-^ (i.e. eGFP^-^) cells and conducted scRNAseq of the two populations using Smart-seq2 protocol (Awasthi et al., 2009; Picelli et al., 2013). Notably, we added IL-23 to the cytokine combination in this experiment in order to enhance signaling through IL-23R and thus better identify its role in Th1 cells. (2) Secondly, we differentiated naïve T cells from IL-23R-deficient mice (*Il23r^eGFP/eGFP^*) with the same set of cytokines and sorted eGFP^+^ vs. eGFP^-^ cells for the expression analysis by scRNAseq (Figure 1C).

We analyzed the expression of signature Th1 and Th17 cell genes in these *in vitro* differentiated cells and confirmed at the single-cell level that IFN-γ-expressing Th1 cells co-express *Il23r* but do not express Th17 cell signature genes such as *Il17a*, *Il17f* and *Ccr6*. In addition to *Ifng*, the cells prominently expressed Th1 signature genes including *Tbx21* and *Cxcr3* (Figures 1D and S2). A low frequency of cells expressed *Rorc*, encoding the transcription factor RORγt associated with Th17 cell differentiation, but the expression of *Rorc* was minimal when compared to the Th1 cell master transcription factor *Tbx21* (Figures 1D and S2).

Comparing the IL-23R^+^ vs. the IL-23R^-^ subsets from the wildtype cells (*Il23r^eGFP/wt^*) provided a way to identify genes that may be co-expressed with IL-23R (Figures 1E and F). Here, we identified a set of 528 differentially expressed (DE) genes whose expression in Th1 cells is positively or negatively associated with the presence of IL-23R and may be involved in pathogenicity of Th1 cells similar to how IL-23R signaling promotes pathogenicity of Th17 cells (Lee et al., 2012a; McGeachy et al., 2009). Indeed, the set of genes that were co-expressed with IL-23R included critical effector molecules and transcription factors including *Il18r1* and *Stat5a* (Figure 1E and table S1).

Strikingly, we found that the number of differentially expressed genes between the eGFP^+^ and eGFP^-^ populations in knockout cells (*Il23r^eGFP/eGFP^*) was substantially lower than the IL-23R^+^ vs. IL-23R^-^ comparison in wildtype cells (129 vs 528 at FDR < 0.1 with only 33 genes overlapping). These eGFP^+^ knockout cells would reflect differentiation in response to IL-12 plus IL-21 in the absence of IL-23R signaling. This finding suggests that the majority of genes we identified in the wildtype cells as being differentially expressed between the IL-23R^+^ and IL-23R^-^ populations are more likely to be directly affected by IL-23R signaling.

We then undertook a combined analysis of the expression profiles from both wildtype and IL23R-deficient cells to better distinguish genes directly regulated by IL-23R signaling in Th1 cells (Figure 1G). To this end, we combined the datasets and incorporated model terms for the presence of eGFP, the genotype (wildtype or IL-23R-deficient), and the interaction of eGFP and genotype (see methods for full details). By testing the interaction term, we sought genes whose expression difference between eGFP^+^ and eGFP^-^ cells was dependent on the genotype. In this manner we identified altogether 219 genes where the presence of wildtype IL-23R was associated with an increase in expression in eGFP^+^ cells compared to eGFP^-^ cells. Of those, *Dusp2*, *Gpr18*, *Trim27*, *Itgb1*, *Gzmb*, *Pde4b*, *Plek*, *Thap11*, *Panx1*, *Peli1*, *Rac1* and *Lamp1*(CD107a) appeared of particular interest as novel candidate genes dependent on IL-23R signaling and potentially important for pathogenicity of Th1 cells (Figure 1G and table S1). Some of these genes have been implicated previously in autoimmune inflammation, such as *Panx1* (Velasquez et al., 2016) and *Rac1* (Kurdi et al., 2016). Our finding that these genes are regulated in an IL-23R-dependent manner suggests an important function in Th1 cells.

In summary, in addition to pathogenic Th17 cells, we find that Th1 cells can be induced to express IL-23 receptor *in vitro* through culture with IL-12 + IL-21 and that the single-cell transcriptomic profile corresponds to that of Th1 cells. Through this process, we have further identified a unique gene set associated with IL-23R signaling in Th1 cells.

### IL-23R^+^ Th1 and IL-23R^+^ Th17 cells share common gene signatures but also display subset-specific features

To investigate whether IL-23R^+^ Th1 and IL-23R^+^ Th17 cells share common features, we differentiated pathogenic Th17 cells with IL-1β + IL-6 + IL-23 and sorted IL-23R^+^ (i.e. eGFP^+^) and IL-23R^-^ (i.e. eGFP^-^) cells for single-cell analysis comparing their transcriptional profiles with those of Th1 cells described above. Interestingly, we find a set of 147 genes whose expression in IL-23R^+^ and IL-23R^-^ cells is shared by Th1 and Th17 cells (Figure 2A and table S1). Among the genes highly expressed in both IL-23R^+^ Th1 and Th17 cells, we find *Il22*, which has been identified previously as a core component of the pathogenicity signature of Th17 cells (Gaublomme et al., 2015; Lee et al., 2012b). Some genes (179) were expressed only in Th1 cells differentially between IL-23R^+^ and IL-23R^-^ cells such as *Il18r1* and *Ifng* (Figure 2B and table S1). Interestingly, 93 genes showed the exact opposite profile in Th1 and Th17 cells, one of which was *Il27ra* which was highly expressed in IL-23R^-^ Th1 cells (Figure 2C and table S1). Finally, *Il17a* and *Il17f* were prominently expressed in IL-23R^+^ cells in Th17 cells only, and were among 158 genes that were differentially expressed between IL-23R^+^ cells and IL-23R^-^ cells only in Th17 cells (Figure 2D and table S1). These results suggest that IL-23R induces a common Th1/Th17 pathogenicity expression profile but also induces a distinct set of genes in Th1 and Th17 cells that likely contribute to their pathogenicity to exert lineage-specific phenotype and function.

**Figure 2.**
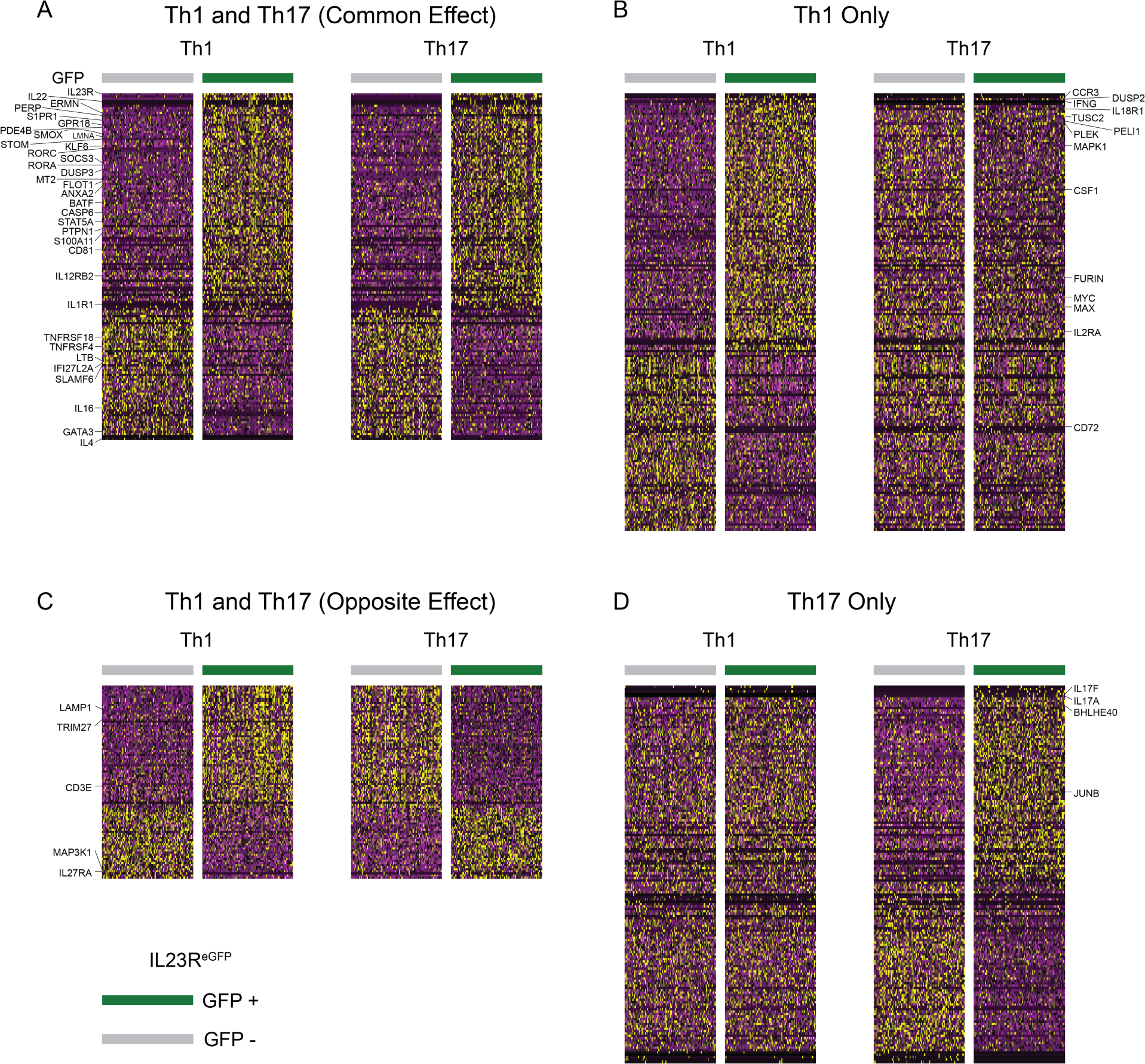
IL-23R^+^ Th1 and IL-23R^+^ Th17 cells show both common and unique transcriptional signatures. Pathogenic Th17 cells were differentiated with IL-1β + IL-6 + IL-23 and IL-23R^+^ (i.e. eGFP^+^) and IL-23R^-^ (i.e. eGFP^-^) were sorted and then single-cell Smart-seq2 was performed and the transcriptional signatures were compared to Th1 cells differentiated with IL-12 + IL-21 + IL-23 reported in Fig. 1. (A) Th1 cells and Th17 cells show a set of genes that is similarly regulated in both IL-23R^+^ cells from either population. (B) Some genes are differentially regulated between IL-23R^+^ and IL-23R^-^ cells in Th1 cells only. (C) Some genes show the exact opposite behavior in Th1 and Th17 cells. (D) A fourth set of genes was differentially regulated between IL-23R^+^ and IL-23R^-^ cells in Th17 cells only.

### IL-23R deficiency protects from Th1 cell adoptive transfer colitis

GWAS analysis has implicated *IL-23R* as a major risk gene for IBD (Duerr et al., 2006), but the actual mechanism by which IL-23R signaling promotes the disease has not been fully elucidated. It has been known that naïve T cells that lack IL-23R lose the ability to induce adoptive transfer colitis, presumably by not developing Th17 cells and promoting differentiation of FoxP3^+^ Tregs (Ahern et al., 2010; Izcue et al., 2008; Powrie et al., 1994). However, as *bona fide* Th1 cells elicit adoptive transfer colitis (Harbour et al., 2015), we assessed the issue of whether IL-23R has a role in these pathogenic Th1 cells to induce colitis. To test the contribution of IL-23R to the pathogenicity of Th1 cells *in vivo*, we differentiated wildtype (*Il23r^eGFP/wt^*) and IL-23R-knockout (*Il23r ^eGFP/eGFP^*) Th1 cells *in vitro* with IL-12 + IL-21 + IL-23 (see methods for details) and then adoptively transferred them to RAG1^-/-^ recipients (Figure 3A). Strikingly, we found that Th1 cells that lack IL-23R do not efficiently induce severe colitis, as determined by the histopathological exam of the colon (Figure 3B). Our results therefore establish an important function of IL-23R in the pathogenicity of Th1 cells and their ability to induce colitis.

**Figure 3.**
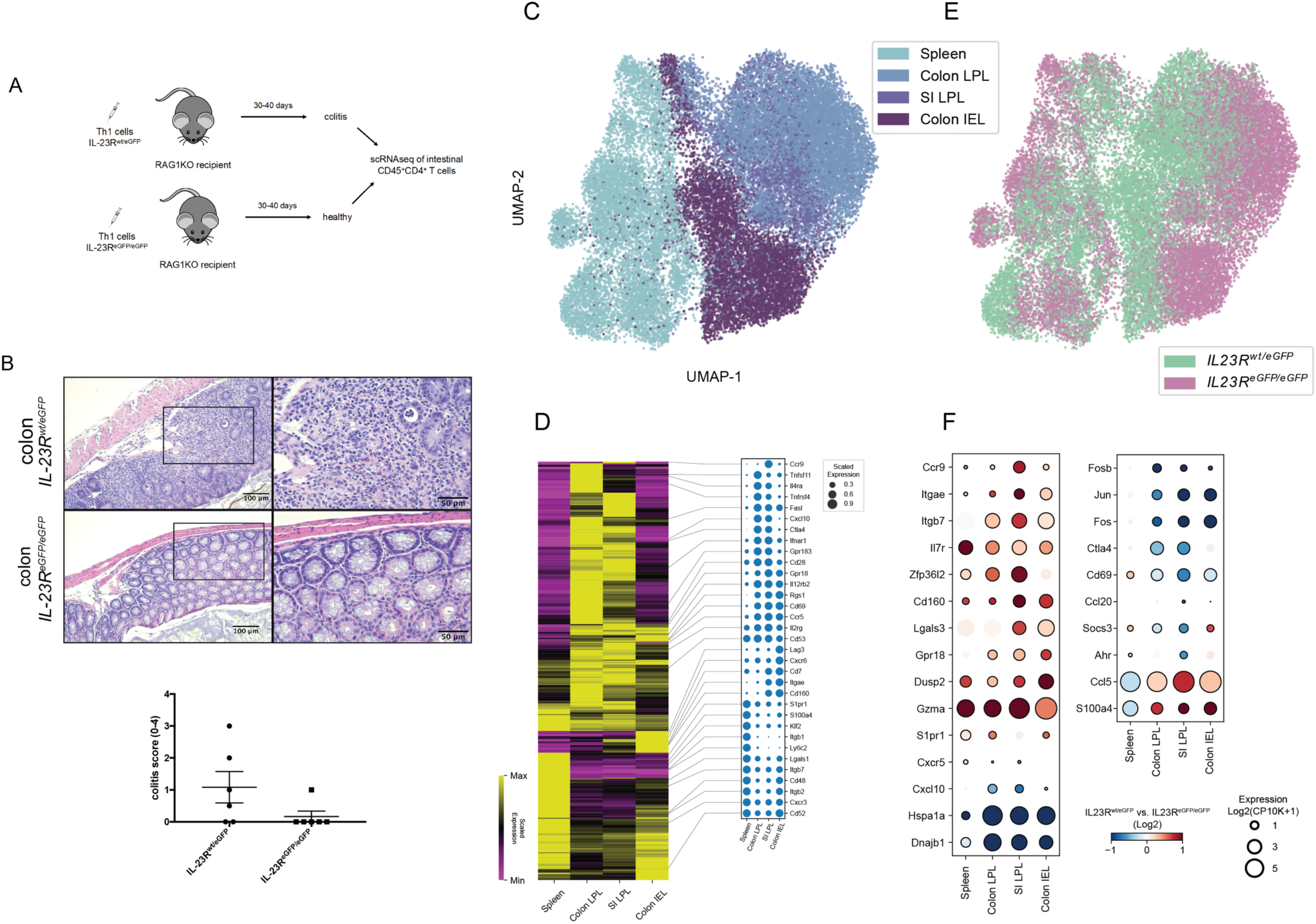
scRNAseq of tissue-infiltrating Th1 cells in a pre-clinical model of adoptive transfer colitis. (A) Schematic of adoptive transfer colitis by *in vitro* differentiated Th1 cells followed by histopathology and 10x scRNAseq. Viable CD45^+^CD4^+^ T lymphocytes from RAG1^-^/^-^ recipients of either wildtype cells (*Il23r^wt^/^eGFP^*) or knockout cells (*Il23r^eGFP^/^eGFP^*), respectively, were isolated from the intestinal mucosa through FACS in preparation of scRNAseq. (B) Deficiency of IL-23R protects from Th1 cell mediated adoptive transfer colitis as evaluated by histopathology. H&E staining and clinical score are shown. (C) UMAP of 30,260 single cells sequenced from spleen, colonic and small intestinal (SI) lamina propria (LPL) and colonic epithelium (IEL = intraepithelial lymphocytes). (D) Heatmap of differentially expressed genes among the tissues analyzed with selected genes highlighted as dot plots. (E) Identification of wildtype (teal) and knockout cells (magenta), respectively. (F) Selected genes are shown that are expressed in an IL-23R dependent manner. Red representing higher expression in wildtype cells and blue representing higher expression in knockout cells.

To test whether the transferred Th1 cells produced IFN-γ or had transdifferentiated into IL-17A-producing Th17 cells, we isolated colonic intraepithelial lymphocytes (IEL) as well as lamina propria lymphocytes (LPL) and tested these cells for the production of IFN-γ, IL-17A and GM-CSF. Animals that received wildtype (*Il23r^eGFP/wt^*) Th1 cells had a dramatically increased frequency of CD4^+^ intraepithelial lymphocytes and lamina propria lymphocytes, up to 60-70% of the entire CD45^+^ population vs. 4%-9% in recipients of knockout (*Il23r^eGFP/eGFP^*) Th1 cells, pointing to the ongoing inflammatory process (Figures S3A, D and F). Interestingly, we found that in the animals studied here, the CD4^+^ T cells had largely maintained their production of IFN-γ and had not significantly deviated into IL-17A-producing cells (Figures S3B, E and G). Furthermore, a sizable frequency of IFN-γ-producing cells in the colon also produced GM-CSF (Figure S3C). It is known that both in the EAE model and the colitis model we used here, Th17 cells show extensive plasticity and become IL-17A^+^/IFN-γ^+^ double positive (Harbour et al., 2015; Hirota et al., 2011). Our results suggest that at least in our settings, Th1 cells were not plastic and retained a Th1 cell phenotype. Taken together, these data suggest that colitogenic Th1 cells require IL-23R expression in order to induce disease *in vivo* similar to Th17 cells and that these cells do not transdifferentiate towards a Th17 phenotype. Therefore, pathogenicity mediated by IL-23R signaling is separable and not limited to Th17 cells *in vivo*.

### Single-cell RNA-sequencing reveals transcriptional signatures of colitogenic intestinal lymphocytes and their dependence on IL-23R signaling

To gain a better understanding of the transcriptional programs that may be driven by IL-23R in transferred Th1 cells and therefore associated with pathogenicity, we isolated CD45^+^CD4^+^ infiltrating T cells from the intestinal mucosa (both the intraepithelial lymphocytes, IEL, and the lamina propria lymphocytes, LPL) from animals that received either wildtype or IL-23R KO cells by fluorescence-activated cell sorting (FACS). We then performed massively parallel single-cell transcriptional profiling by 10x technology of 32,763 cells (Figure 3A).

The cells primarily cluster depending on which tissue they were isolated from, with splenic CD4^+^ T cells segregating from cells of intestinal origin (Figure 3C). These results confirm previous findings that T cells acquire distinct transcriptional profiles depending on whether they are isolated from secondary lymphoid organs such as the spleen or reside in tissues such as the intestine (DiSpirito et al., 2018). We find the expression of 1224 genes was strongly associated with the tissue of origin (FDR < 0.1, |logFC| > 0.5), whereas other genes showed a broader expression in cells of all tissue origins (Figure 3D and table S2). For instance, CD69 has been proposed as a marker of tissue residency in T cells including in T_RM_ (tissue resident memory) cells and the precise tissue specific expression of *Rgs1* suggests that it could be involved in trafficking and tissue residency as well, in particular in the intestinal mucosa (Masopust and Soerens, 2019). Klf2, a transcription factor that is important for the expression of CD62L and S1pr1, and which was shown to direct lymphocyte trafficking was highly upregulated in splenic cells suggesting an antagonistic function to CD69 and potentially Rgs1 as well (Carlson et al., 2006). Interestingly, the chemokine *Cxcl10* was particularly highly expressed in T cells within the LPL but not IEL (Figure 3D), suggesting a very specific autocrine/paracrine regulation, in particular since the contribution of *Cxcl10* to the trafficking and retention of pro-inflammatory Th1 cells expressing the corresponding receptor *Cxcr3* has been previously identified (Bonecchi et al., 1998; Singh et al., 2003). Overall, multiple chemokine-related transcripts were expressed in a compartment-specific fashion and therefore likely contribute to the trafficking of Th1 cells to the intestine (Figure 3D).

In terms of their splenic transcriptional profiles, it appeared that the wildtype Th1 cells (*Il23r^eGFP/wt^*) and the knockout cells (*Il23r^eGFP/eGFP^*) isolated from recipient mice were rather uniformly distributed (Figure 3E). Strikingly, the wildtype and KO cells appeared to instead segregate within the intestinal tissues, suggesting that Th1 cells from the wildtype vs. IL-23R KO mice may attain transcriptionally different expression profiles when they infiltrate the intestine (Figure 3E). We then asked which genes would be differentially expressed between wildtype and KO cells across the four analyzed tissues (spleen, LPL small intestine (SI), LPL colon and IEL colon) and identified between 84 and 190 genes per tissue (FDR < .1 and |logFC| > 0.5) (table S2). We identified genes such as *Gpr18*, *Cd160* and *Zfp36l2* being differentially expressed dependent on IL-23R signaling in all or most tissues *in vivo* (Figure 3F). Interestingly, the chemokine Ccl5 (RANTES), a ligand of Ccr5, exhibited a clear IL-23R dependent expression within the intestine only, suggesting that the regulation of Ccl5 is particularly relevant within the tissue and that IL-23R might contribute to pathogenicity by enabling inflammatory cells to migrate to tissues. *Cxcl10* did not show such IL-23R-dependent regulation (Figure 3F). Taken together, our observations support the notion that IL-23R signaling affects the tissue-specific transcriptional expression of novel genes relevant to intestinal inflammation.

### IL-23R drives the expansion of highly inflammatory and colitogenic Th1 cells in the lamina propria as uncovered by scRNAseq

Lymphocytes residing within the lamina propria exert a critical contribution to elicit intestinal inflammation (Khor et al., 2011). Therefore, we decided to focus on these samples (both from the colon and small intestine) in wildtype (*Il23r^eGFP/wt^*) and IL-23R KO (*Il23r^eGFP/eGFP^*) cells (Figure 4A). We partitioned all cells into clusters and identified a total of 12 distinct clusters with characteristic transcriptional profiles within the LPL (Figure 4B). We then sought to identify clusters consisting of cells with a highly pro-inflammatory and colitogenic gene signature that may be responsible for mediating colitis. Importantly, we noted differential contribution of wildtype or KO cells to some clusters, especially clusters 2, 8, 9 and 10 were dominated by cells from wildtype animals (*Il23r^eGFP/wt^*) whereas clusters 5 and 7 were dominated by KO cells (*Il23r^eGPF/eGFP^*) (Figure 4C). Based on their transcriptional profiles, we found that clusters 2 and 9 consisted of cells that exhibited a highly colitogenic gene signature and may have the ability to drive intestinal inflammation which will be highlighted in detail below (Figure 4D).

**Figure 4.**
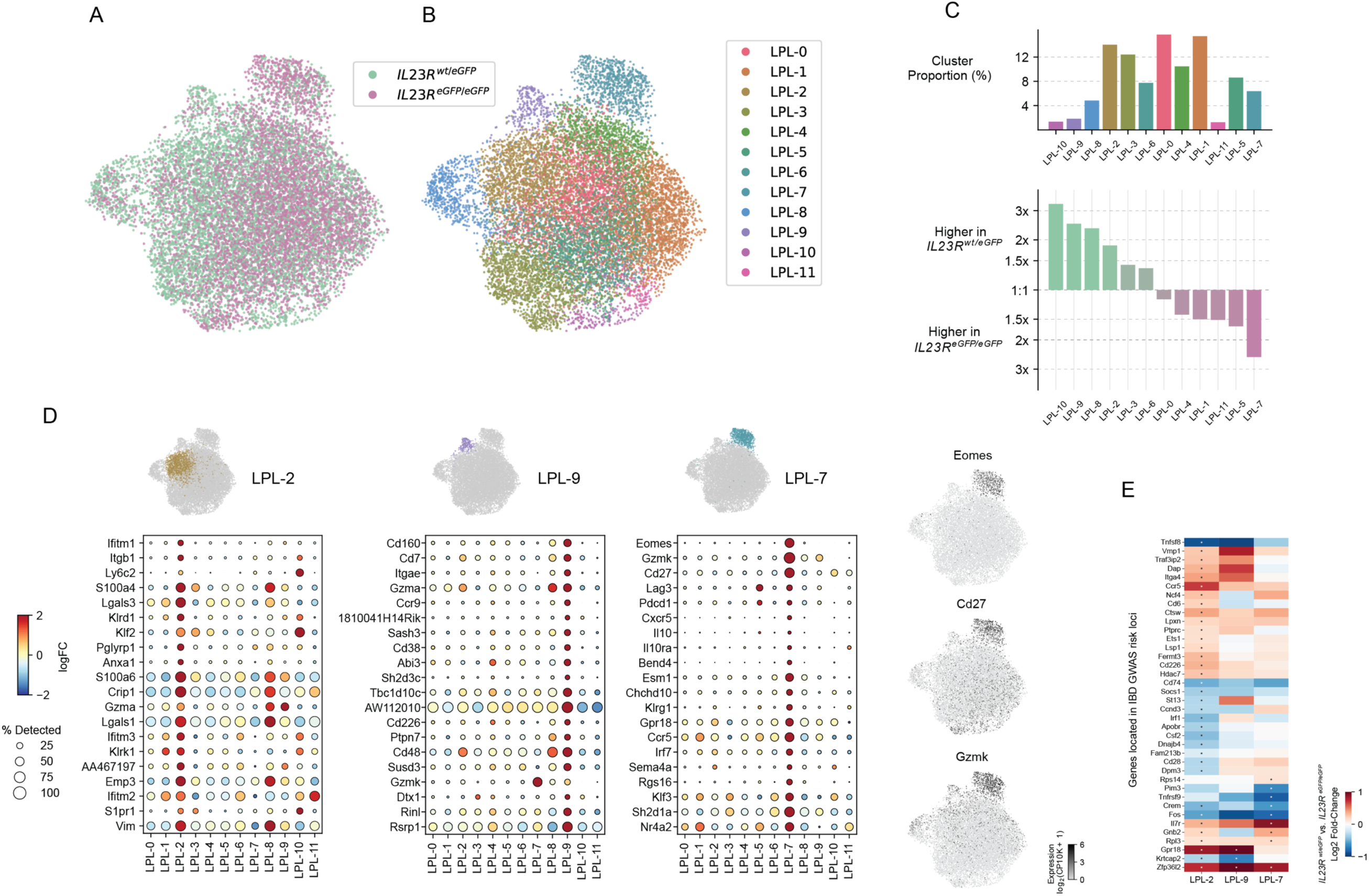
IL-23R drives the expansion of highly inflammatory and colitogenic T cells in the lamina propria as identified by scRNAseq. (A) UMAP of T cells isolated from the lamina propria of small intestine and colon. Wildtype cells (teal) and knockout cells (magenta) are highlighted. (B) Cluster analysis identifies 12 clusters with particular transcriptional signatures. (C) The size of each cluster is shown as a percentage of the total number of cells. For each cluster, its relative abundance among wildtype vs. knockout cells is visualized. (D) Two clusters with a highly inflammatory and colitogenic signature are shown (clusters 2 and 9) which are dominated by wildtype cells. The top 20 differentially expressed genes in comparison to all other clusters are shown. Cluster 7 consists in the majority of knockout cells and exhibits a signature reminiscent of Tr1-like cells. (E) We assembled a list of genes (597 genes) found within IBD GWAS risk loci for which De Lange *et al*. (2017) provided the main basis. We then asked which of these genes is expressed in clusters 2, 7 and 9 in an IL-23R-dependent manner comparing expression levels between wildtype and knockout cells. Positive values indicate higher expression in wildtype cells (*Il23r^eGFP/wt^*). Asterisks indicate statistically significant differences (FDR<0.1).

We found that the characterizing genes of cluster 9 (one vs. all comparison, FDR < 0.1, |logFC| > 0.5 and table S3) encode a wide range of molecules essential for inflammation and colitis. For example, among the most highly expressed genes was *Cd103/Itgae* which has been shown to enable pathogenic T cells to migrate to their target tissue and is characteristic of pro-inflammatory colonic T cells in ulcerative colitis (Annacker et al., 2005; Lamb et al., 2017). Another highly expressed gene, *Ccr9*, encodes a gut homing receptor and has been shown to play a crucial role in colitis (Johansson-Lindbom and Agace, 2007). Previously, it was shown that the transmembrane glycoprotein CD38 is important for intestinal inflammation (Schneider et al., 2015). Likewise, CD48, a SLAM (signaling lymphocyte activation molecule) family member, is crucial for T cell function in colitis and it has been suggested that CD48 may represent a useful target in human IBD (Abadia-Molina et al., 2006). Strikingly, an ORF (open reading frame) within the gene *AW112010* (encoding a long-noncoding RNA) which was among the top 20 differentially regulated genes within cluster 9 in our data set has been recently shown to be essential in mucosal immunity including colitis (Jackson et al., 2018). The expression of this ORF is fairly broad across clusters and the locus appears to be transcribed in the vast majority of cells. However, the expression of *AW112010* shows a dramatic increase in expression within the inflammatory cluster 9 (Figure 4D).

Importantly, in addition to the genes in cluster 9 that have been implicated in the development of colitis, the cluster also contained novel genes not previously associated with the development of colitis. For example, *Cd160*, the most upregulated gene in cluster 9 encodes an Ig superfamily member (Anumanthan et al., 1998) and has not been assigned a function in colitogenic T cells yet. The gene encoding the nectin receptor family member *Cd226* (Dnam-1), was also part of the signature of highly expressed genes in cluster 9. Co-stimulatory receptors such as CD226 play a critical role in T cell activation, autoimmunity and cancer (Zhang and Vignali, 2016). Previously, CD226 was shown to be crucial for the activation of cytotoxic lymphocytes and Th1 cells and its expression within cluster 9 suggests that it may play an important role in intestinal inflammation (Dardalhon et al., 2005; Gilfillan et al., 2008). In summary, the transcriptional analysis of cluster 9 exhibits an entire array of genes critical for intestinal inflammation and identifies novel potential genes important for intestinal inflammation such as *Cd160*.

Many genes highly expressed in cluster 2 are implicated in IFN-γ signaling (Figure 4D and table S3). In fact, IFN-γ signaling has early been identified as being critical for intestinal inflammation in various pre-clinical models of colitis (Ito et al., 2006; Singh et al., 2003). For example, top differentially genes in this cluster were *Ifitm1* (interferon-inducible transmembrane protein), *Ifitm2* and *Ifitm3*. Members of this family are interferon response genes that have been shown to be entry sites in viral infection (Brass et al., 2009). A study in human IBD patients identified IFITM1 as a potential prognostic marker in ulcerative colitis (Roman et al., 2013). Interestingly, we identified many integrins as being part of the list of genes highly expressed within cluster 2, with *Itgb1* as the second highest differentially expressed gene and which has previously been implicated in colorectal cancer (Laudato et al., 2017). Two additional integrins highly upregulated in cluster 2 (albeit not among the top 20 differentially expressed genes) were *Itga4* (position 101) and *Itgb7* (position 32). Strikingly, a recent study demonstrated that targeting the leukocyte integrin *α*_4_β_7_ with vedolizumab was more effective in moderately to severely active ulcerative colitis than adalimumab (a humanized monoclonal antibody neutralizing TNF) (Sands et al., 2019a).

The observation, that many genes implicated in IFN-γ signaling and lymphocyte trafficking (integrins) were highly upregulated in cluster 2 suggests that this cluster consisted of inflammatory cells that could potentially represent the cellular phenotype of early potential drivers of the intestinal inflammatory response.

The other two clusters that were enriched in wildtype cells (*Il23r^eGFP/wt^*) were clusters 8 and 10.

Cluster 8 represented a cluster of highly proliferating cells exemplified by the expression of Ki-67 and genes important for cell cycle progression such as *Cdc* (cell division cycle) genes, including *Cdc6* (critical for the initiation of DNA synthesis (Mailand and Diffley, 2005)) and *Cdc20*, which is known to activate the anaphase promoting complex/cyclosome APC/C (Yu, 2007) (Figure S4). The observation that cluster 8 was highly enriched with wildtype cells suggests that these cells were actively proliferating within the tissue to contribute to the inflammatory process.

Cluster 10 on the other hand showed high expression of *Ccr7* which is commonly correlated with an ability of T cell trafficking and homing to lymph nodes and Peyer’s patches (Forster et al., 2008). In addition, cluster 10 cells expressed higher levels of the anti-apoptotic gene *bcl2* which is known to be driven by STAT5 signaling and may be an interesting target in IBD as suggested in pre-clinical studies (Weder et al., 2018).

In summary, using single-cell RNA-sequencing, we identify distinct clusters of T cells within the intestinal mucosa that exhibit a highly inflammatory profile, in particular clusters 2 and 9. Furthermore, the observation that these clusters are dominated by wildtype cells suggests that IL-23R signaling critically contributes to the observed expression of inflammatory genes. Importantly, we succeed in identifying novel genes such as *Cd160* that may play an important role for intestinal inflammation and IBD.

### IL-23R is implicated in the reciprocal regulation of Tr-1 like cells and limits their expansion

Cluster 7 caught our particular attention as it is strongly enriched with IL-23R KO cells which would suggest that this cluster contains predominantly non-pathogenic T cells (Figure 4C). The 117 genes that characterize this cluster (one vs. all comparison, FDR < 0.1, logFC > 0.5) include *Eomes*, *CD27*, *Gzmk*, *Lag3* and *Il10* (Figures 4D, S5A and S5B). This finding was of particular significance as these genes have been recently identified to define a novel human IFN-γ^+^IL-10^+^ Tr1-like cell type that may be reduced in IBD patients (Gruarin et al., 2019). In fact, Eomesodermin (Eomes) has been shown to be a key transcription factor of a Tr1-like lineage (Zhang et al., 2017). Our finding that this cluster 7 is predominantly composed of IL-23R KO cells suggests that IL-23R may function as an important gatekeeper and negative regulator for the development of Tr1-like cells in the colonic mucosa. Previously, it was shown that IL-23R is indeed expressed on these IFN-γ^+^IL-10^+^ Tr1-like cells and therefore IL-23 signaling could potentially regulate development and function of this population (Alfen et al., 2018). To our knowledge, our study is the first to provide functional evidence that IL-23R may act as a negative regulator of such Tr1-like cells. In addition, PD1 was shown to be expressed on human Tr1-like cells which is highly consistent with our data presented here (Alfen et al., 2018) (Fig. 4D). In fact, it was shown that PD1 is implicated in the conversion of human T-BET^+^ Th1 cells to FOXP3^+^ Tregs (Amarnath et al., 2011). Other co-inhibitory molecules such as Lag-3 were also more highly expressed in cluster 7 (Figures 4D and S5B). We used a published transcriptional signature that characterizes human Tr-1 like cells and were able to show that it identifies cluster 7 (upregulated compared to other clusters, Wilcoxon rank-sum test p < 10^-200^) providing strong evidence that the cells of cluster 7 are the equivalent of these human cells in our study (Figure S5C) (Gruarin et al., 2019). In summary, our results suggest that IL-23R signaling balances the differentiation of colitogenic Th1 vs. Tr1 cells and therefore effector vs regulatory arms of the immune response.

Interestingly, comparing WT to KO cells within cluster 7 revealed a set of heat shock protein family members, in particular *Hspa1a* and *Hspa1b* (Figure S5D). Heat shock proteins are implicated in the regulation of T cell responses and specific single nucleotide polymorphisms have recently been identified in HSPA1L that distinguish IBD patients from healthy controls (Takahashi et al., 2017; van Eden et al., 2005). Our finding suggests that heat shock proteins of the Hsp70 family, in particular Hspa1a and Hspa1b, may be protective in IBD and regulated in an IL-23R dependent manner.

### Comparative analysis of human IBD GWAS with our scRNAseq study nominates genes of particular interest for the function of colitogenic Th1 cells in an IL-23R dependent manner

It has become clear from genome-wide association studies (GWAS) that IL-23R plays an important role in IBD (Duerr et al., 2006). To investigate if IL-23R signaling is particularly relevant to the expression of other genes found within human IBD GWAS loci in T cells, we conducted a survey of the roughly 240 IBD risk loci that have been identified so far (de Lange et al., 2017; Huang et al., 2017; Jostins et al., 2012; Liu et al., 2015) and tallied their associated genes (about 700 genes in total)(de Lange et al., 2017). Within loci that contain multiple genes, we focused on genes that were found to be implicated, for example, by fine-mapping or by expressed quantitative trait loci (eQTL) analysis which reduced the list of genes that we further investigated to 597. Since we established that clusters 2 and 9 were dominated by wildtype cells with a highly inflammatory transcriptional signature we focused the analysis on these two clusters in addition to cluster 7 which was dominated by KO cells to investigate the potential contribution of genes found in GWAS loci to IL-23R mediated pathogenicity. We used within-cluster comparison of WT vs. KO cells and identified genes found within IBD GWAS loci that were expressed in an IL-23R dependent manner (Figure 4E and table S3). This finding suggests that IL-23R signaling in Th1 cells may be relevant to induction of genes implicated through GWAS in the development of IBD.

Among the genes in the set implicated through GWAS were *Gpr18*, *Traf3ip2*, *Ncf4* and *Ets1*.

*Gpr18* was particularly highly expressed in clusters 2 and 9 in an IL-23R dependent manner. Recently, it was shown that certain mutations are increased in ulcerative colitis including in the gene *Traf3ip2* (encoding Act1) which is part of the NFKBIZ pathway (Kakiuchi et al., 2020). *Ncf4* plays an important role in cellular reactive oxygen species (ROS) pathways as part of the NOX2 (NADPH oxidase 2) complex. Ets1 is a transcription factor and essential co-factor of Tbet in Th1 cell-mediated inflammatory responses (Grenningloh et al., 2005).

Adhesion and migration determine major biological outcomes of T cell-mediated autoimmunity. Interestingly, a number of genes implicated in these processes were part of the identified gene signature of cluster 2, in particular *Lsp1*, *Lpxn* (Leupaxin) and *Fermt3*. We hypothesize that IL-23R signaling may exert pathogenicity also through downstream effects on migration and adhesion.

Notably, some genes were not limited to clusters 2 and 9 in their IL-23R-dependent expression such as *Zfp36l2* but exhibited this pattern also across other clusters including cluster 7 suggesting that their genotype-dependent expression may be too dominant than to be confined to a particular cluster (Figure 4E). Zfp36l2 belongs to a family of RNA binding proteins that have been implicated in Th17 cell biology but their contribution to Th1 cell-mediated autoimmunity remains unknown (Gaublomme et al., 2015; Wells et al., 2017).

Our findings illuminate genes implicated in cellular pathways ranging from trafficking, cellular adhesion (integrins), metabolic regulators and immune cell interaction identified in GWAS IBD risk loci that may prove critical in their contribution to intestinal inflammation driven by IL-23R in colitogenic T cells. The observation that these genes found in GWAS IBD risk loci are driven by IL-23R in colitogenic T cells may suggest that IL-23R and these genes converge mechanistically in a disease-relevant manner.

### Ranking identifies novel genes as drivers of intestinal inflammation in an IL-23R dependent manner

In many cases, the combinatorial analysis of several data sets yields a comprehensive insight into useful targets for further analysis as critical genes may not only emerge as being the top differentially expressed genes in a singular data set. Therefore, we combined both *in vitro* and *in vivo* single-cell RNA-sequencing data sets we reported above and included data from human IBD GWAS to rank the genes that may have an impact on the development of colitis.

We postulated the following criteria that were important to us and queried our data set for genes that would match as many criteria as possible: (1) As we were particularly interested in genes potentially conferring pathogenicity, we focused on genes that were most highly expressed in clusters 2 and 9 in comparison to all other clusters and that were differentially expressed between wildtype and KO cells within either cluster 2 or 9; (2) Genes of particular importance to intestinal inflammation may exhibit a highly tissue specific expression and therefore we evaluated their expression comparing splenic and intestinal samples; (3) We asked which genes showed an IL-23R-dependent expression in Th1 cells differentiated under IL-12 + IL21 + IL-23 *in vitro*; (4) We incorporated as one criterion whether a given gene was previously identified in human GWAS to confer risk and/or is located within a GWAS risk locus for IBD (Figure 5A and table S4, see methods for details of ranking).

**Figure 5.**
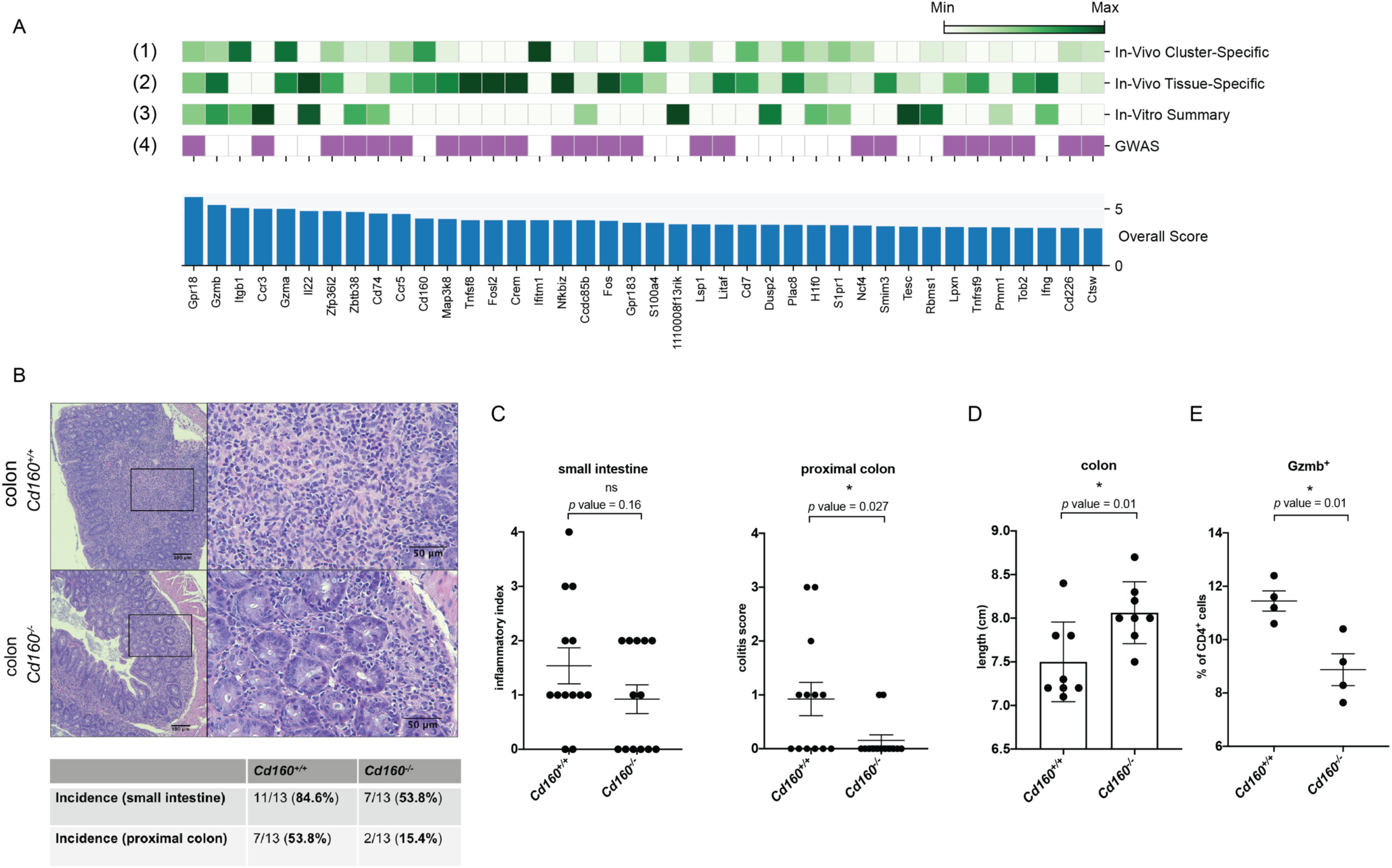
Ranking algorithm and validation identify novel drivers of T cell-mediated intestinal inflammation in an IL-23R dependent manner. (A) Identification and ranking of potential novel drivers of intestinal inflammation taking several critical considerations into account. 4 tracks are shown. Track 1: Genes are evaluated based on their expression *in vivo* in a cluster and genotype specific manner emphasizing clusters 2 and 9. Track 2: Genes are ranked based on their tissue specific expression comparing intestinal and peripheral (splenic) expression. Track 3: Genes are ranked based on their IL-23R dependent expression *in vitro* following differentiation with IL-12 + IL-21 + IL-23 (Smart-seq2 data from Fig. 1). Track 4: Genes that are found within an IBD GWAS risk locus are identified contributing to an increased ranking score. See methods for details of ranking. (B) Deficiency for CD160 in Th1 cells adoptively transferred into RAG1^-/-^ recipients protects from colitis. Representative H&E histopathological stainings are shown. (C) Intestinal inflammation and colitis are scored in the following way: 0 (healthy) – 4 (most severe colitis). Pooled data from three independent experiments are shown for small intestine and proximal colon. Mean + s.e.m. are indicated. (D) Colon length in RAG1^-/-^ recipients of either wildtype or CD160^-/-^ cells. Pooled data are shown from two independent experiments. (E) Gzmb expression is measured by flow cytometry in both Cd160^+/+^ and Cd160^-/-^ Th1 cells prior to adoptive transfer.

*Gpr18* scored very highly in all criteria including being located within a GWAS risk locus (Jostins et al., 2012). Interestingly, it was shown that Gpr18 plays a crucial role for CD8aa^+^ intraepithelial T lymphocytes within the intestinal mucosa (Wang et al., 2014). Our studies suggest it may be a critical mediator of pathogenicity by IL-23R in colitogenic Th1 cells. The transcription factor NF-*κ*B is critical to T cell function and in particular mutations within the NFKBIZ pathway were recently found to be relevant in ulcerative colitis (Kakiuchi et al., 2020). *Nfkbiz* itself scored highly in our ranking data as a target in colitogenic T cells (Figure 5A).

### CD160 plays an important role in colitogenic Th1 cells

Finally, we sought to provide conclusive evidence of the critical contribution of candidate genes identified in our studies to T cell-driven intestinal inflammation. To this end, we acquired knockout mice for *Cd160*, the top differentially expressed signature gene of inflammatory cluster 9 and which ranked prominently in the ranking scheme (Figure 5A) (Tan et al., 2018). CD160 is an Ig superfamily member and its function in inducing colitogenic T cells and IBD has not been investigated. It has been shown to be important for IFN-γ production in NK cells and being expressed on intraepithelial CD8^+^ T cells (Tan et al., 2018; Tu et al., 2015). We differentiated naïve T cells from these CD160 KO mice to Th1 cells *in vitro* as described above and tested their ability to elicit adoptive transfer colitis. We did not observe a difference between wildtype and CD160 KO Th1 cells to produce IFN-γ or GM-CSF prior to adoptive transfer *in vitro* (Figures S6A and B). The cells did not express any IL-17A prior to adoptive transfer. Strikingly, the recipient animals that received CD160 deficient Th1 cells showed a strong protection from colonic inflammation as assessed by histopathological examination (Figure 5B). The decrease of inflammation showed a clear trend in the small intestine and was statistically significant in the colon (Figures 5B and C) The histopathological analysis was consistent with the shortening of the colon in recipients of wildtype cells (Figure 5D). When we isolated T cells from mesenteric lymph nodes from recipients of either wildtype or CD160 knockout cells we did not observe a significant difference in IFN-γ expression which was consistent with our observations *in vitro* suggesting that it was likely not any lack in the ability to produce IFN-γ that protected recipients of CD160 KO cells from developing severe colitis (data not shown). Previously, it was shown that CD160 is important for *Gzmb* expression in intestinal CD8^+^ T cells which helps to clear Listeria monocytogenes infection (Tan et al., 2018). Interestingly, this was highly consistent with our observation that CD160 KO cells differentiated *in vitro* with IL-12 + IL+21 + IL23 produced less Gzmb compared to wildtype cells which could contribute mechanistically to their diminished colitogenicity *in vivo* (Figure 5E). Taken together, our results demonstrate an important function of CD160 to the colitogenicity of Th1 cells in an IL-23R dependent manner.

## Discussion

IL-23R function has been considered a critical driver of pathogenicity in Th17 cells and important for the regulation of FoxP3^+^ Tregs but not for other T helper cell subsets. However, several experimental and clinical observations have been difficult to reconcile under this assumption. We provide compelling evidence that IL-23R plays an important role in other pathogenic T cell subsets, in particular a subset of IL-23R^+^ Th1 cells.

First, we report that *bona fide* Th1 cells can be differentiated *in vitro* with IL-12 + IL-21 and express similar levels of IL-23R as pathogenic Th17 cells polarized with IL-1β + IL-6 + IL-23. This illustrates how the combinatorial action of cytokines crucially determines the functional outcome since IL-21 has been shown to participate with TGF-β to induce pro-inflammatory Th17 cells, not Th1 cells (Korn et al., 2007; Nurieva et al., 2007). In Th17 cells, IL-6 has been shown to induce IL-21 that promotes IL-23R expression in an autocrine manner, and this induction is STAT3 dependent (Zhou et al., 2007). Which transcription factor mediates the induction of IL-23R in Th1 cells differentiated with IL-12 + IL-21 remains to be determined. However, one such transcription factor could be c-MAF that has been shown to bind to the IL-23R promoter and to work in conjunction with IL-21 in some non-Th17 cells, in particular in the setting of autoimmune diabetes which is known to be Th1 cell-driven (Hsu et al., 2018; Iwamoto et al., 2014; Sato et al., 2011).

We continue to show that IL-23R is required in Th1 cells *in vivo* to acquire full pathogenicity to elicit adoptive transfer colitis. We take advantage of scRNAseq to profile 32,763 cells in a pre-clinical model of colitis including roughly 20000 intestinal T cells and uncover transcriptional signatures that are governed by IL-23R and that are critical to the emergence of pathogenic and colitogenic Th1 cells. The inflammatory signatures of our pre-clinical model were highly reminiscent of human studies in IBD as outlined above. We identify novel candidate genes for T cell-mediated IL-23R dependent pathology and provide a framework of genes identified in previous IBD GWAS as potential drivers in colitogenic Th1 cells. Our study therefore contributes to the important effort of identifying and validating genes found within GWAS risk loci having an important function in colonic inflammation and their implication in colitogenic T cells.

Previously, it was shown that Th17 cells show extensive plasticity during an inflammatory response and that Th17 cells eventually may develop features of Th1 cells by upregulating *Tbx21* and expressing IFN-γ both in mouse and human (Acosta-Rodriguez et al., 2007; Becattini et al., 2015; Hirota et al., 2011). IL-23R was shown to play an important role in this lineage plasticity (Hirota et al., 2011) and it was assumed that IL-23R bearing Th cells that express IFN-γ are solely transdifferentiated Th17 cells. Interestingly, it was shown in the adoptive transfer model of colitis that Th17 cells convert to Th1 cells and that this appears to be essential to Th17 cell-mediated disease in this model (Harbour et al., 2015). Importantly, we show that *bona fide* Th1 cells can be differentiated to express *Il23r in vitro* and that IL-23R deficient Th1 cells lose their ability to elicit intestinal inflammation. Interestingly, we did not observe extended plasticity in the Th1 cells adoptively transferred in our study towards a Th17 cell phenotype. With the exception of cluster 3 in the LPL that contained cells showing a transcriptional signature of Th17 cells including expression of *Ccr6*, *Il17a*, *Il22*, *Il17re* and *Il17f*, however, we did not identify IL-17A^+^ cells by intracellular cytokine staining when retrieved from the intestinal mucosa in RAG1 KO recipients. Thus, the vast majority of LPL cells showed a strong transcriptional signature of *bona fide* Th1 cells including expression of *Il12rb1*, *Il12rb2*, *Cxcr3*, *Stat1*, *Stat4*, *Tbx21* and *Ifng*.

We speculate that Th1 cells represent a further advanced state of inflammatory trajectory that may be reached with or without transitioning through a Th17 cell state. Strikingly, it has been demonstrated that both Th17 and Th1 cells may ultimately give rise to T cells with regulatory function at the resolution of inflammation (Amarnath et al., 2011; Gagliani et al., 2015).

Importantly, we describe here a dichotomous function of IL-23R in promoting inflammatory cells on one hand and on the other hand limiting the expansion of Tr1 cells with a regulatory function, identified by expression of signature genes such as *Eomes*, *CD27*, *Il10ra*, *Gmzk* and co-inhibitory receptors *Lag-3* and *Pdcd1*. A concept is evolving in which IL-23R serves not only as a critical driver of inflammatory T cells but a central integrator of immune responses within the local immunological milieu. In fact, in addition to the emergence of Tr1-like cells in IL-23R deficient cells in our study, it has been shown previously that IL-23R is also expressed on colonic FoxP3^+^ Tregs, making these cells receptive to sudden inflammatory changes that limit their suppressive effector functions in the presence of the pro-inflammatory cytokine IL-23 (Izcue et al., 2008; Schiering et al., 2014).

In the setting of Th17 cells, it was shown that the salt-sensing kinase Sgk1 is important in regulating their plasticity toward FoxP3^+^ T cells (Wu et al., 2018). Which mediators downstream of IL-23R signaling suppress a Tr1 cell fate in Th1 cells remains to be determined.

A ranking scheme combining both *in vitro* and *in vivo* transcriptional signatures and their dependence on IL-23R signaling enabled us to identify novel drivers of T cell pathogenicity and we succeeded in validating one of these candidates, CD160. Indeed, CD160 emerged in a study profiling the cellular composition and transcriptional signatures of ileal biopsies from Crohn’s patients although the functional significance remained unclear (Uniken Venema et al., 2019). Interestingly, CD160 was shown to be very important for intraepithelial type I ILCs which are amplified in Crohn’s disease and was important for the pathogenicity of ILC1s in the anti-CD40 colitis model in addition to control of L. monocytogenes infection. These data suggest that CD160 is not only important for CD4^+^ T cells as demonstrated in our study but for other immune cells such as ILC1s and CD8^+^ T cells as well (Fuchs et al., 2013; Tan et al., 2018). To validate that we had not inadvertently included immune cells other than CD3^+^CD4^+^ T cells in our analysis that have been shown to express CD160, we confirmed that the cells sequenced lacked expression of NK cell markers and expression of CD8 (Figure S7).

Another important candidate that emerged and which scored highly in all categories in the ranking (Figure 5A) was Gpr18. Interestingly it was identified as a central part of a transcriptional signature in myeloid cells driving a Th1 cell intestinal inflammation triggered by the colonization of germ-free mice with Klebsilla isolated from a Crohn’s patient (Atarashi et al., 2017)

In summary, we uncouple IL-23R as a purely Th17 cell-specific pathogenicity factor and implicate it in being critical for evoking a pathogenic phenotype in Th1 cells. Furthermore, through the powerful combination of a highly relevant pre-clinical model of colitis with massively parallel single-cell RNA-sequencing and integration of GWAS data from IBD patients, we identify novel potential drivers of IL-23R-mediated T cell pathogenicity and validate one of these candidates. We hope that the discovery of the role of IL-23 in the induction of pathogenic Th1 cells and their contribution to autoimmune disease on one hand and the inhibition of IL-10 producing Tr1 cells on the other hand may identify novel targets and prove useful for the development of therapeutic intervention in multiple human autoimmune diseases. Indeed, our findings provide a reason to revisit the pathogenic function of IL-23R in autoimmune diseases in which Th1 or other T cell subsets are believed to be the main drivers of the disease.

## Acknowledgments

We would like to thank the Kuchroo and Anderson laboratories for feedback, in particular Davide Mangani. Furthermore, we would like to thank Mary Collins for feedback on both manuscript and data. We thank Arlene Sharpe (HMS) for generous sharing of CD160 KO animals. We would like to thank the Harvard rodent histopathology core for help with histology and analyses. We would like to acknowledge the flow cytometry core facility at the Ann Romney Center for Neurologic Diseases (ARCND) for cell sorting, in particular Deneen Kozoriz and Rajesh K. Krishnan. Funding: This study was supported by grants 5P01AI073748-10, 1R01AI144166-01A1, 5P01AI056299-17, 5P01AI039671-23 (V.K.K.) and the Chan Zuckerberg Biohub (N.Y.). C.W. is supported by a National Multiple Sclerosis Society grant (TA-1605-08590).

## Author contributions

M.P., G.M.z.H., Y.L., D.D, N.Y. and V.K.K. conceived the study. M.P. performed most functional studies with preliminary data generated by G.M.z.H. and Y.L. J.N. and D.D. helped with single-cell sequencing. C.W., A.W., P.R.B. and A.C.A. provided critical feedback and discussion. Computational analyses of single-cell data were performed by D.D. and N.Y. with additional help from S.J.R. M.P., D.D., N.Y. and V.K.K. wrote the manuscript with help from G.M.z.H and R.J.X. The study was supervised by N.Y and V.K.K. with additional guidance by A.R. and R.J.X.

## Declaration of interests

N.Y. is an advisor and/or has equity in Cellarity, Celsius Therapeutics and Rheos Medicines. R.J.X. is cofounder and equity holder in Celsius Therapeutics and Jnana Therapeutics. A.C.A. is a member of the scientific advisory boards for Compass Therapeutics, Tizona Therapeutics, Zumutor Biologics, and ImmuneOncia, which have interests in cancer immunotherapy. A.C.A. and V.K.K. are paid consultants for iTeos Therapeutics. V.K.K. has an ownership interest in and is a member of the scientific advisory board for Tizona Therapeutics. V.K.K. is a co-founder of and has an ownership interest in Celsius Therapeutics. V.K.K is a co-founder of Bicara therapeutics. A.C.A.’s and V.K.K.’s interests were reviewed and managed by the Brigham and Women’s Hospital and Partners Healthcare in accordance with their conflict of interest policies.

---

**Figure S1.**
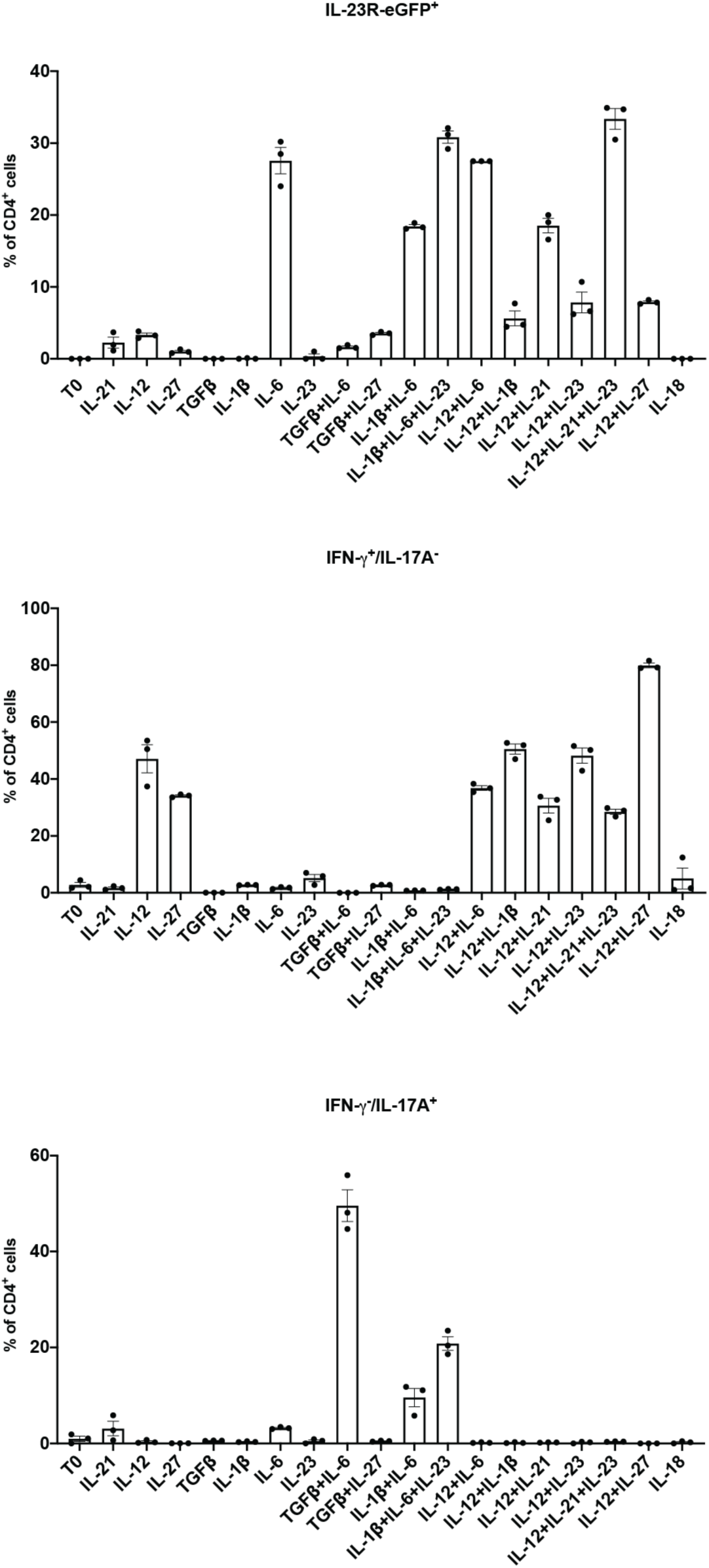
Cytokine screen identifies IL-21 as a cytokine that together with IL-12 induces strong expression of IL-23R in Th1 cells *in vitro*. We isolated naïve T cells from *Il23r^wt/eGFP^* reporter mice and tested 19 different conditions. IL-23R expression was measured by flow cytometry identifying eGFP^+^ cells. Several conditions strongly induced IL-23R expression including the pathogenic Th17 cell condition IL-1β + IL-6 + IL-23. The strongest expression of IL-17 was observed with the condition TGF-β + IL-6. Even though IL-21 and IL-6 alone induced IL-23R expression, they resulted in the complete lack of IFN-γ expression and showed slight induction of IL-17A. Importantly, both IL-12 + IL-21 and IL-12 + IL-21 + IL-23 induced strong expression of the Th1 cell signature cytokine IFN-γ and complete lack of IL-17A expression.

**Figure S2.**
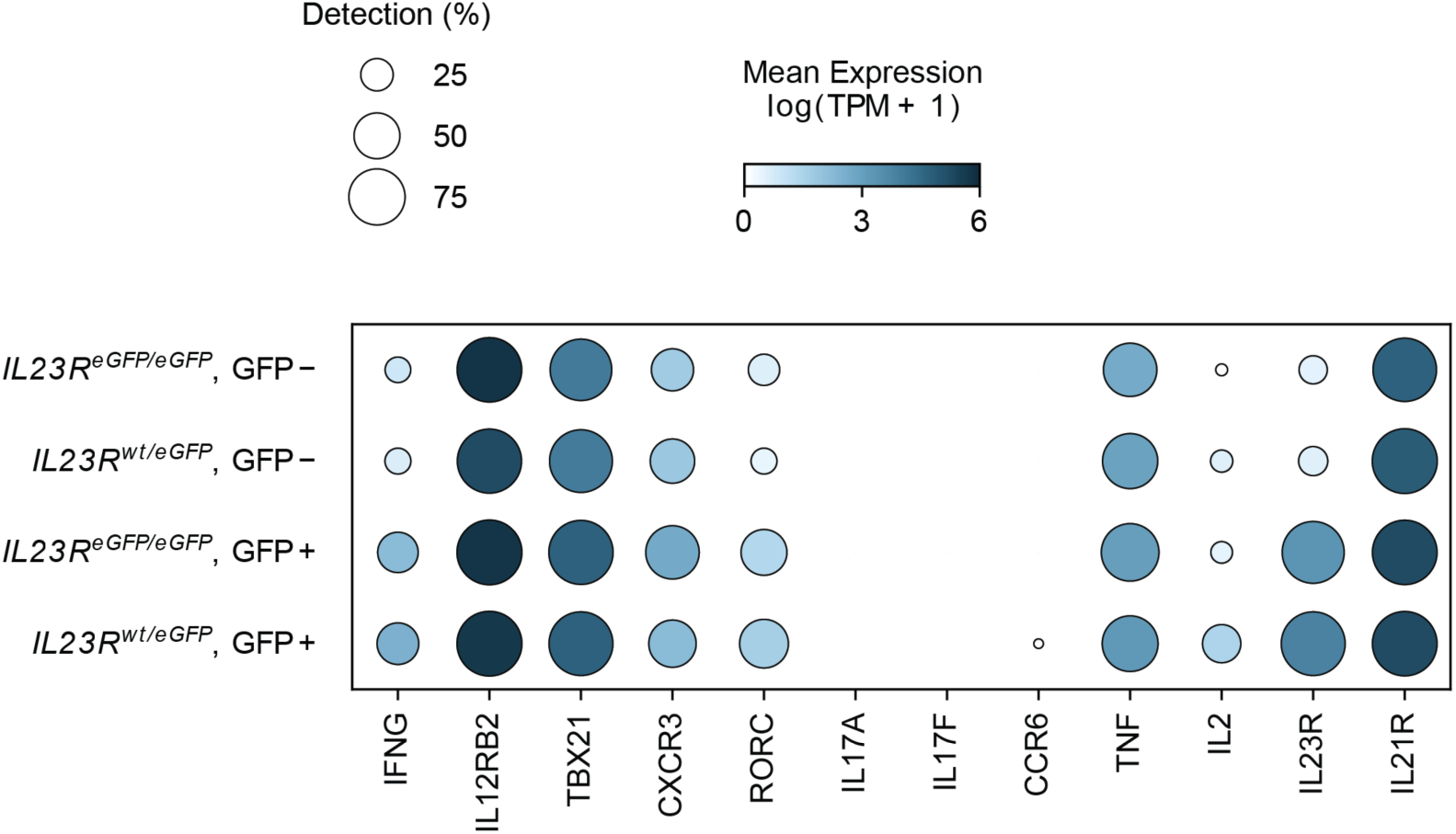
Dot plot showing expression of 12 key genes in Th1 and Th17 cell biology across the 4 populations profiled by Smart-seq2. *Tbx21*, *Il12rb2* and *Il21r* expression appeared largely unaltered by deficiency for *Il23r* which was consistent with unaltered expression between eGFP^+^ and eGFP^-^ populations. Of note, the expression level of *Il23r* in knockout cells (*Il23r^eGFP/eGFP^*) is merely reminiscent of the fact that these cells produce mRNA truncated of the essential C-terminal region of IL-23R which was replaced by an *IRES*-*eGFP* sequence which results in a functional KO but is detected by Smart-seq2.

**Figure S3.**
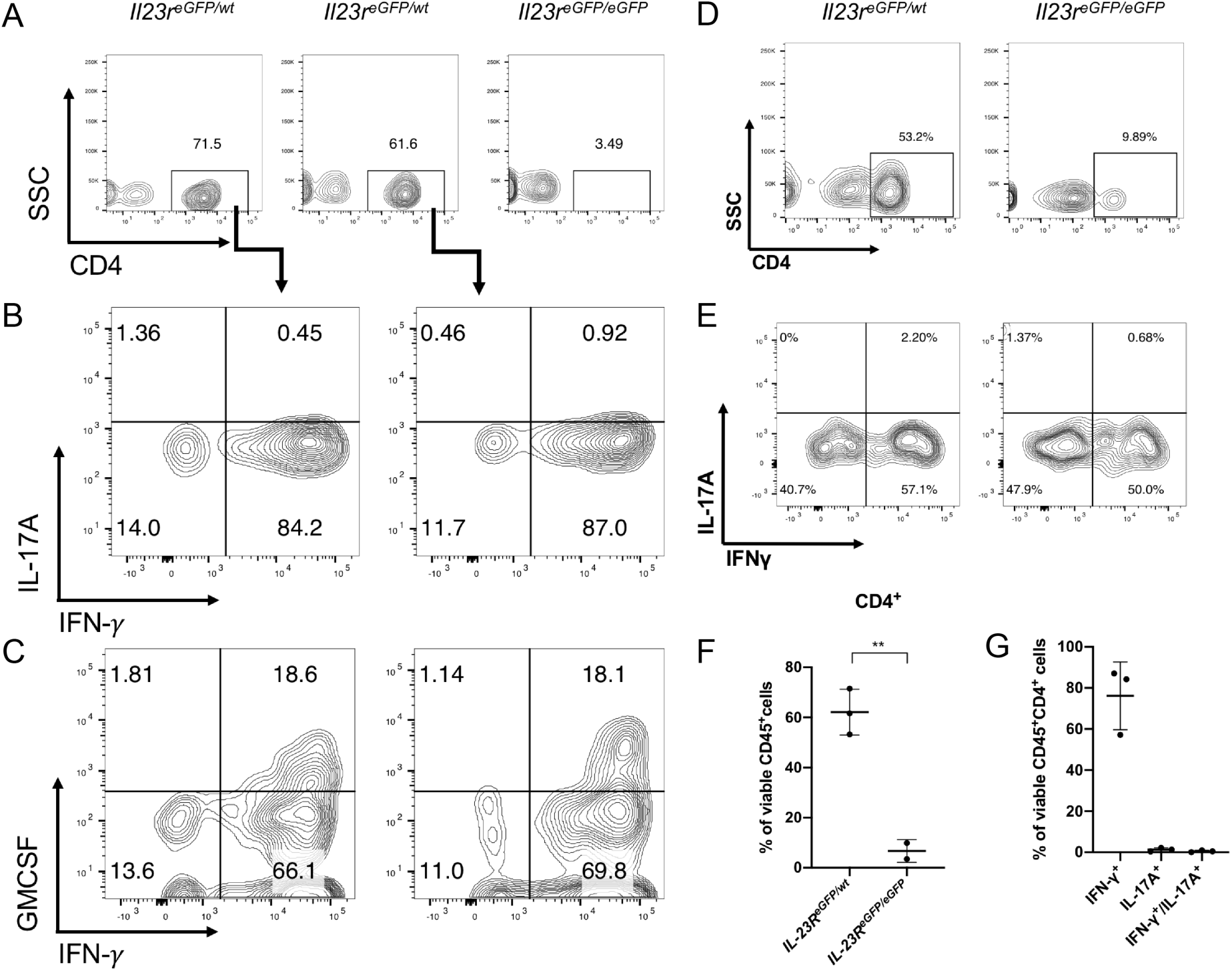
Tissue infiltrating CD45^+^CD4^+^ T cells show strong expression of IFN-γ. Flow cytometric analyses of tissue infiltrating T cells isolated from the intestine of recipients of either wildtype or IL-23R KO cells and cultured ON with the cytokines IL-7+IL-23. ICC was performed after restimulation with PMA/ionomycin. (A) Colonic LPL cells. Two recipients of wildtype cells (*Il23r^eGFP/wt^*) and one recipient of KO cells (*Il23r^eGFP/eGFP^*) are shown. Gated on viable CD45^+^ cells. (B) and (C) ICC for IFN-γ, IL-17A and GMCSF of the two recipients of wildtype cells is shown. (D) Colonic IEL cells. One recipient of wildtype cells and one recipient of KO cells are shown. Gated on viable CD45^+^ cells. (E) ICC for IFN-γ and IL-17A of the IEL samples. (F) and (G) Pooled values of LPL and IEL samples. LPL and IEL samples were isolated in independent experiments and combined for analysis. Data in panels (F) and (G) are mean ± SD. Unpaired t-test, p value ** < 0.01 in panel (F).

**Figure S4.**
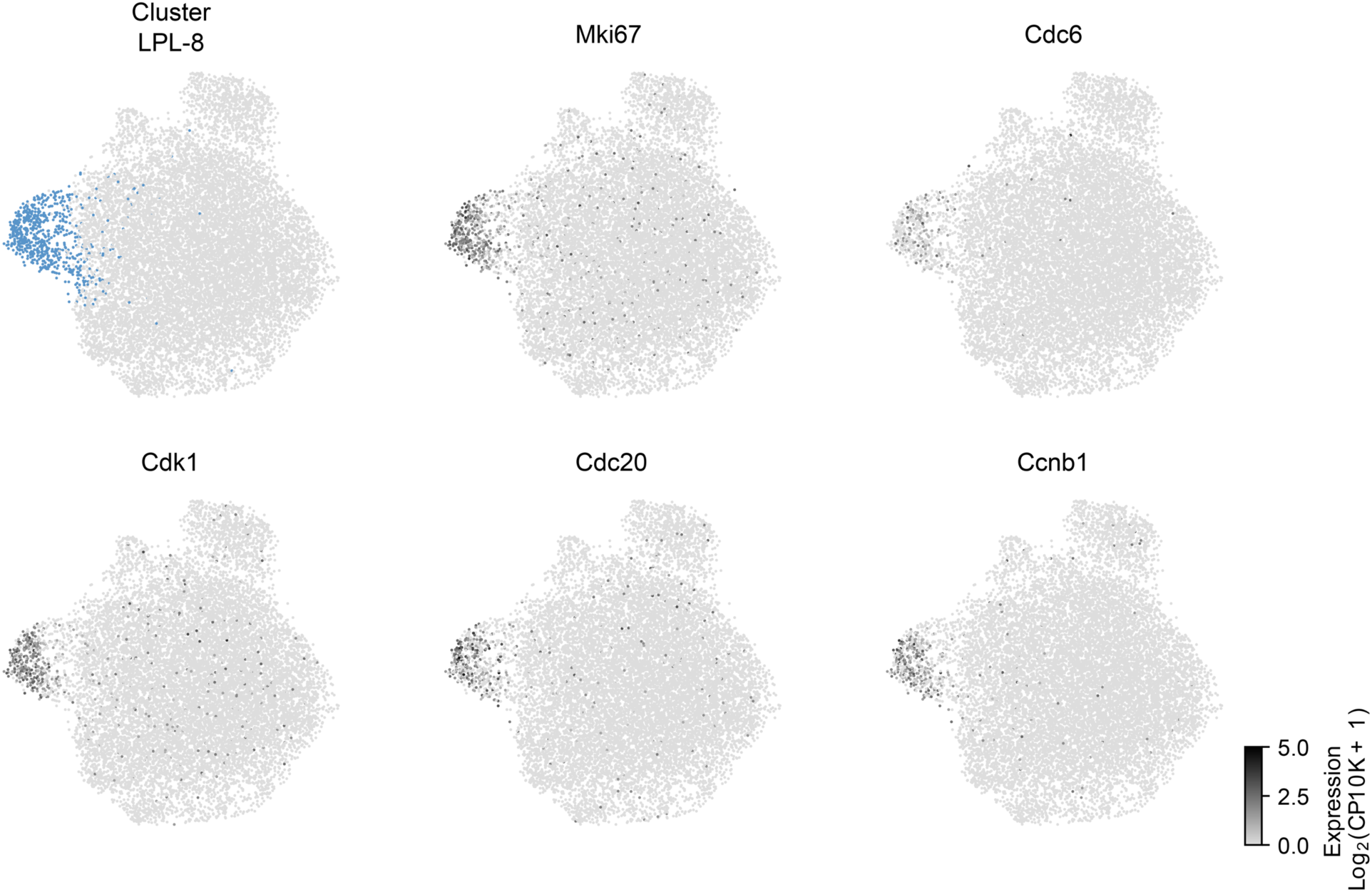
Cluster 8 represents highly proliferating cells. UMAPs show that cells of cluster 8 highly express genes critical to cell cycle progression such as *Cdc20*, *Ccnb1* (cyclin B), *Cdc6*, *Cdk1* and the proliferative marker *Mki67*.

**Figure S5.**
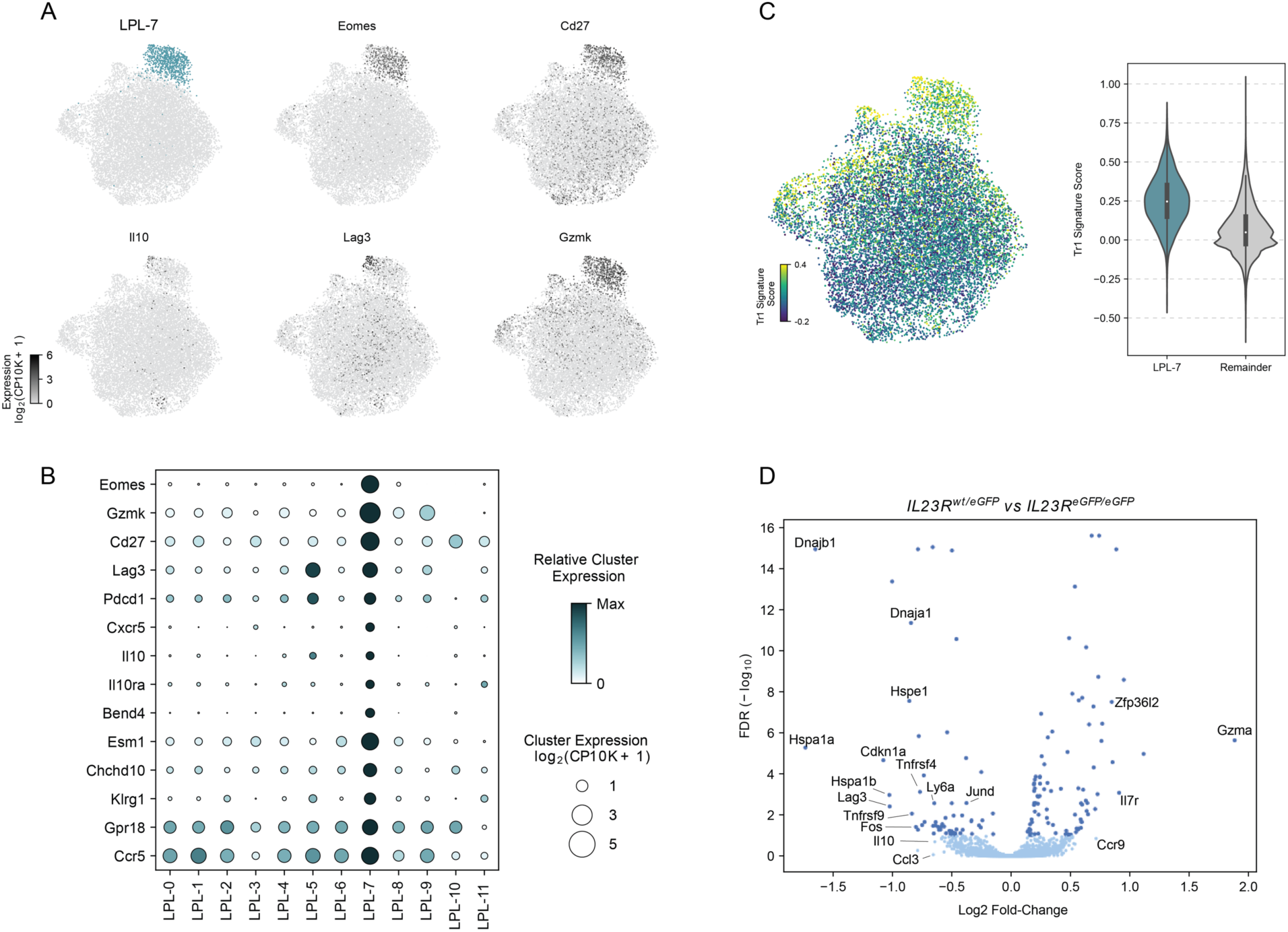
IL-23R negatively impacts the development of regulatory Tr1 cells in the intestinal lamina propria as identified by scRNAseq. (A) UMAPs and correlated expression profiles of signature genes of regulatory Tr1 cells exposing cluster 7. (B) Dot plot showing the expression of selected signature genes in comparison to all other clusters. (C) Transcriptional signature of Tr1 regulatory cells identified by Gruarin *et al*. (2019) highlights cluster 7. (D) Volcano plot highlighting differentially expressed genes between wildtype (*Il23r^eGFP/wt^*) and knockout (*Il23r ^eGFP/eGFP^*) cells within cluster 7.

**Figure S6.**
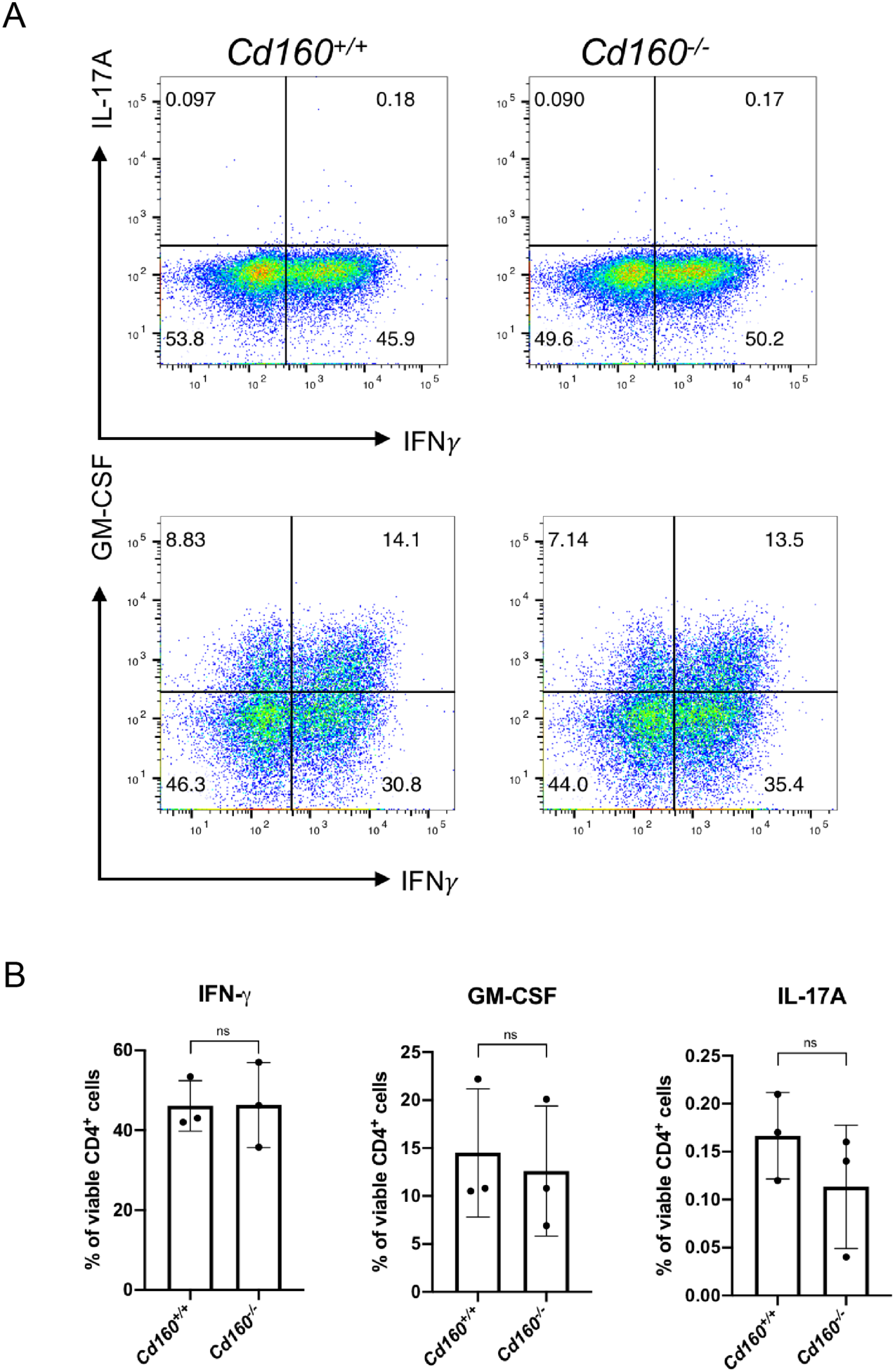
*In vitro* differentiated Th1 cells from either *Cd160^+/+^* or *Cd160^-/-^* cells do not show a difference in IFN-γ expression prior to adoptive transfer. (A) *In vitro* differentiated Th1 cells do not show a difference in IFN-γ production under culture with IL-12 + IL-21 + IL-23 between wildtype and CD160 knockout cells. One representative experiment of three independent experiments is shown. (B) Pooled data from *in vitro* differentiated Th1 cells are shown prior to adoptive transfer into RAG1^-/-^ mice. 3 independent experiments. Mean is shown with SD error bars. All differences are non-significant.

**Figure S7.**
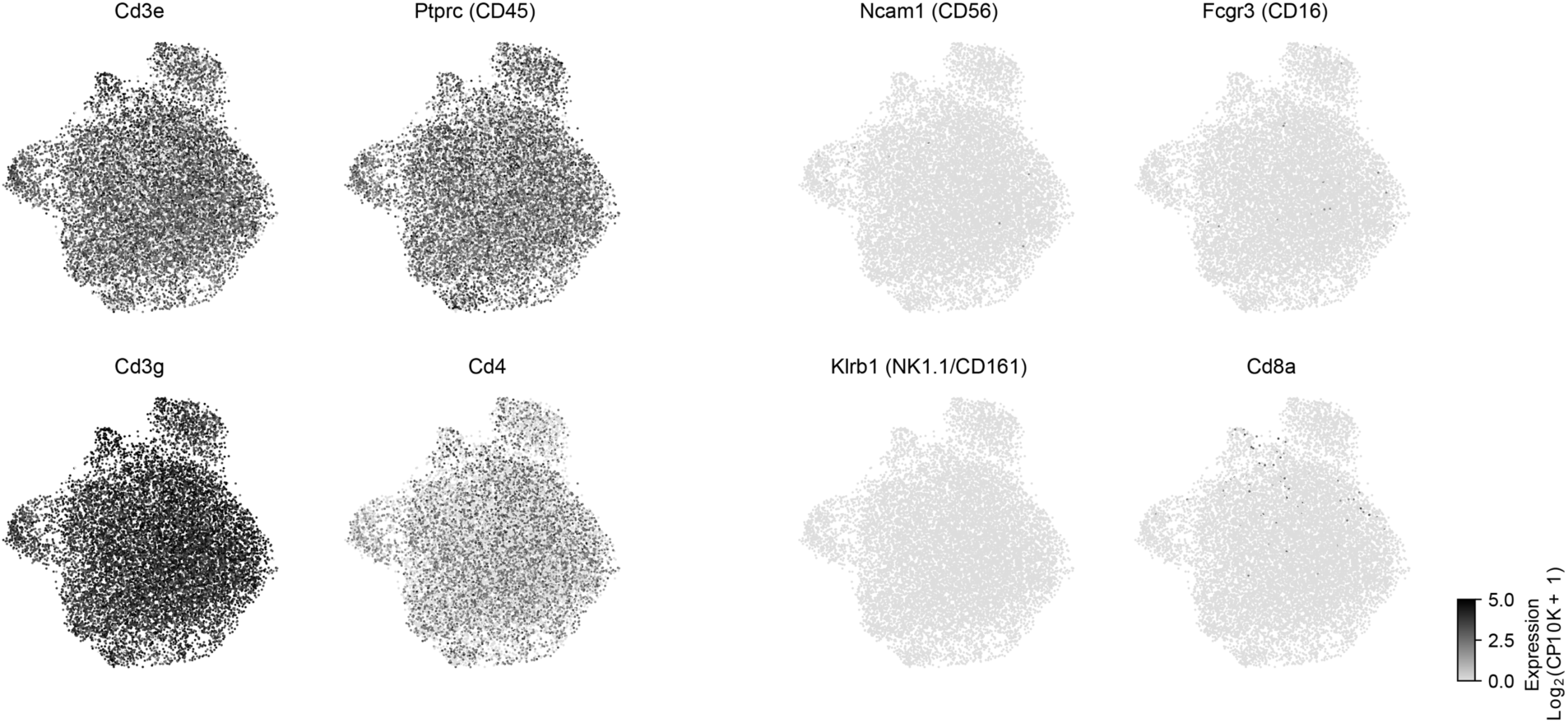
Overview of T cell and NK cell markers in cells isolated from the intestinal mucosa (LPL) and sequenced by 10x technology. UMAPs of cells sequenced that show uniform expression of CD45, CD3, CD4 and lack expression of NK cell markers such as NK1.1, CD56 and CD16 in addition to the lack of CD8a.

## Materials and Methods

### Mice

IL-23R^eGFP^ reporter mice were described previously (Awasthi et al., 2009). The following strain was purchased from The Jackson Laboratories: B6.129S7-Rag1^tm1Mom^/J (known as Rag1^-/-^; Stock No.: 002216). CD160^-/-^ mice were a kind gift of Dr. Arlene Sharpe and previously described (Tan *et al*., 2018). All animals were bred and maintained under specific-pathogen-free (SPF) conditions. Genotyping was performed with DNA isolated from tail biopsies. All experiments were approved by and carried out in accordance with guidelines of the Institutional Animal Care and Use Committee (IACUC) at Harvard Medical School and Brigham and Women’s Hospital.

### *In vitro* T cell differentiation

A single-cell suspension was prepared from spleen and lymph nodes and red blood cells were removed by lysis with ACK buffer (Lonza). CD4^+^ T cells were enriched using microbeads (Miltenyi Biotec, Cat. #.: 130-117-043). Naïve CD4^+^ T cells (CD4^+^CD44^low^CD62L^high^CD25^−^) were isolated by fluorescence-activated cell sorting (FACS) using a custom-built BD FACSAria instrument. Sorted naïve CD4^+^ T cells were activated with plate-bound anti-CD3 and anti-CD28 antibodies (1μg/ml; BioXCell; Clone 145-2C11 and clone PV-1, respectively) in 96-well flat-bottom plates (50-100×10^3^ cells/per well). Cells were cultured for 5 days (cytokine screen) with the following cytokine concentrations: IL-1β (20ng/ml); IL-6 (25ng/ml); IL-23 (20ng/ml); IL-12 (20ng/ml); IL-21 (20ng/ml); TGF-β (2ng/ml); IL-18 (20ng/ml). All recombinant cytokines were purchased from Biolegend (IL-21), R&D (IL-12, IL-1β, IL-6, IL-23) and Miltenyi Biotec (hTGF-β).

### Flow cytometry and antibodies

Single-cell suspensions were stained with diluted antibodies in ice-cold PBS + 2% FBS. Dead cells were identified with either 7-AAD (BD Biosciences) or viability dye eFluor 506 (eBioscience). For intracellular cytokine staining (ICC), cells were stimulated for 4 hr at 37°C with phorbol 12-myristate 13-acetate (PMA, Sigma), ionomycin (Sigma) and monensin (GolgiStop; 1:1000; BD Biosciences). Then, cells were stained for surface markers and viability. Cells were fixed and permabilized according to the manufacturer’s instructions (BD Cytofix/Cytoperm kit). All flow cytometric data were acquired with either a BD LSRII or Fortessa instrument and analyzed with FlowJo (BD).

The following antibodies labeled with different fluorophores were purchased from Biolegend unless otherwise indicated: anti-CD4 (RM4-5), anti-CD25 (PC61), anti-CD44 (IM7), anti-CD62L (Mel14), anti-IL-17A (TC11-18H10.1), anti-IFN-γ (XMG1.2), anti-GM-CSF (MP1-22E9, BD), anti-CD45 (30-F11).

### Adoptive transfer colitis

Naïve T cells were isolated from donor mice by FACS and differentiated for 6 days prior to adoptive transfer with the following paradigm. Naïve CD4^+^ T cells (in 6 well dishes, 1×10^6^ cells/ml) were cultured for 48hrs on anti-CD3/anti-CD28 coated plates with the cytokines IL-12 (10ng/ml) + IL-21 (20ng/ml). After 48hrs, the cells were passaged onto non-coated plates and IL-2 (20ng/ml) was added to the culture. Cells were passaged if needed 1:2. On day 4, IL-23 (20ng/ml) was added to the culture. During the entire culture, the medium was supplemented with anti-IL-4 antibody (BioXcell; clone 11B11). On day 5, a small sample was taken and used for ICC to determine IFN-γ^+^ cells and cells to be adoptively transferred were re-stimulated by passage onto anti-CD3/anti-CD28-coated plates for at least 24hrs. After 6 days, cells were harvested, washed in ice-cold PBS and viability was assessed with Typan Blue staining. Cells to be injected were normalized based on IFN-γ expression determined by ICC and 400000 viable IFN-γ^+^ cells were injected intraperitoneally per animal. As recipients, sex-matched RAG1^-/-^ animals were used. The weight was measured pre-adoptive transfer and routinely over the course of the experiment.

For histopathology, the colon and small intestine were fixed in 10% formalin and processed by the Harvard Medical School rodent histopathology core for H&E staining. The severity of disease was assessed in a blinded manner by an experienced histopathologist. The following scoring system was used: 0 (healthy) – 4 (most severe disease).

### Isolation of intestinal lymphocytes

Intestinal lymphocytes were isolated from RAG1^-/-^ recipients using a lamina propria dissociation kit according to the manufacturer’s instructions (Miltenyi Biotec, Cat. No.: 130-097-410). Briefly, intestines were isolated and flushed with ice-cold buffer. The small intestine and colon were separated, opened longitudinally with scissors, turned inside-out and then cut into small pieces (<1cm). Then, the intraepithelial lymphocytes (IEL) were shaken off with buffer containing 1mM DTT with gentle rotation at 37°C. After the incubation period, the IEL were swiftly filtered (100μm), washed with complete medium and kept on ice. This step was repeated once followed by a washing step to remove DTT. Then, the procedure was continued to isolate the lamina propria lymphocytes (LPL). For digestion of the intestinal mucosa, the samples were gently incubated with a proprietary enzymatic solution in C-tubes (Miltenyi Biotec, Cat. No.: 130-096-334) at 37°C for at least 30 minutes using a rotator. After the enzymatic incubation, the samples were homogenized using a gentleMACS (Miltenyi Biotec) and the program m_intestine_01. For overnight culture, to assess the cytokine profile of intestinal T cells, the cells were further isolated using a Percoll (GE Life Sciences/Cytiva) gradient centrifugation to remove cell debris and fat tissue.

### Analysis of *ex vivo* cytokine production by intestinal lymphocytes

IEL and LPL were cultured overnight in complete medium with IL-7 (20ng/ml, Biolegend) + IL-23 (20ng/ml, R&D). The next day, the cells were stimulated with PMA/ionomycin + GolgiStop for 4hrs as described above. ICC was performed and the cells stained for IL-17A, IFN-γ and GM-CSF.

### Single-cell RNA-sequencing (scRNAseq)

For full-length scRNAseq, single, viable *in vitro* differentiated CD4^+^ T cells were sorted into 96-well plates containing 5 μl TCL Buffer (QIAGEN) with 1% 2-mercaptoethanol, centrifuged and frozen at −80 °C. SMART-Seq2 protocol was carried out mainly as previously described (Singer *et al*., 2016). For droplet-based 3′ end massively parallel single-cell RNA-sequencing, sorted (FACS) CD45^+^CD4^+^ intestinal T cells were encapsulated into droplets, and libraries were prepared using Chromium Single Cell 3′ Reagent Kits v2 according to the manufacturer’s protocol (10x Genomics). Generated libraries were sequenced on a HiSeq X (Illumina).

### RNA isolation and quantitative PCR

Total RNA was isolated with the RNeasy kit according to the manufacturer’s instructions (Qiagen). In brief, cells were homogenized with 200μl RLT buffer supplemented with 2-mercaptoethanol by gentle pipetting and then an equal amount of 70% ethanol was added, the samples were mixed by inversion and RNA was captured through a centrifugation step with silica-based columns. Purified RNA was reverse-transcribed using Superscript II enzyme and random hexamer primers (Invitrogen). Taqman probes for genes of interest were purchased from Applied Biosystems. A ViiA7 Real-time PCR system (Applied Biosystems) was used for amplification and data acquisition.

Taqman probes

| Taqman assay | Gene |
| --- | --- |
| Mm00519943_m1 | *Il23r* |
| Mm01168134_m1 | *Ifng* |
| Mm00439618_m1 | *Il17a* |
| Mm00450960_m1 | *Tbx21* |
| Mm01261022_m1 | *Rorc* |
| Mm01219775_m1 | *Stat3* |

### Statistical analysis

Details regarding the statistical analyses can be found in the figure legends. Statistical analysis was performed using GraphPad Prism 7.0 and 8.0. A p-value < 0.05 was considered significant.

### Data availability

All sequencing data that support the findings of this study will be deposited in the Gene Expression Omnibus (GEO).

### Analysis of in-vitro SmartSeq2 single-cell transcriptional profiles

Sequencing reads were initially filtered with Trimmomatic 0.32 (Bolger et al., 2014) to remove low quality ends and adapter sequences. A combination of Bowtie2 v2.2.9 (Langmead and Salzberg, 2012) was used to align reads to a reference transcriptome derived from the GRCm38 Mus musculus genome (NCBI Refseq). RSEM 1.2.31(Li and Dewey, 2011) was used to compile the transcriptome and estimate transcript abundance following read alignment. Prior to alignment, the GRCm38 reference was augmented to include the modified eGFP+IL23R functional knockout transcript.

Prior to downstream analysis, cell libraries were excluded if they contained less than 800,000 reads or more than 3 million reads. Additionally, libraries with less than a 40% alignment rate were withheld. Finally, genes detected in less than 5 cell libraries were discarded prior to differential expression.

Initial inspection of the RNA-seq libraries showed that the dominant source of variation between cells could be attributed to differences in cell cycle progression. To reduce the effect of cell cycle on differential expression testing, two cell-cycle ‘scores’ per cell were computed and input as covariates into the differential expression model. To compute these scores, we first subset the expression matrix, retaining only 97 genes identified as G1/S or G2/M enriched genes in a previous study (Kowalczyk et al., 2015). Then, PCA was run on the log(TPM+1) expression values and the first two component scores per cell were extracted.

To identify differentially expressed genes, the MAST (Finak et al., 2015) R package was used. When fitting the hurdle model, in addition to the covariates of interest both cell cycle components (described above) as well as the number of detected genes per cell were used to reduce false positives due to cycling state or mRNA capture rates. P-values were corrected with the Benjamini Hochberg FDR procedure and genes were accepted at an FDR < 0.1 threshold. To identify genes whose expression differed between the GFP^+^ and GFP^-^ subsets in a genotype-dependent manner (and therefore may be perturbed as a consequence of Il-23 receptor signaling), data from all four subsets (Il23R^WT/eGFP^ and Il23R^eGFP/eGFP^, GFP^+/-^ both) were combined and modeled with categorical covariates for GFP, Genotype, and a GFP:Genotype interaction term (in addition to the cell cycle and detection rate terms). Genes whose expression differed between the FACS sorted GFP subsets were first identified by comparing this full model to a reduced model without the GFP and GFP:Genotype terms, retaining those with FDR < 0.1. Then, in this subset, we tested the GFP:Genotype interaction term specifically and retained genes with FDR < 0.1.

For visualization, the dataset was reduced using principal components analysis (PCA) followed by tSNE (van der Maaten, 2014). Prior to running PCA, the log(x+1) transformed TPM expression values were normalized by regressing out three cell-level covariates: the number of detected genes, and both cell-cycle components. The top 30 components from PCA were then used as input to the tSNE procedure (with perplexity = 30) to further reduce the data to 2 dimensions.

When comparing the GFP-associated expression between T cells stimulated under Th1 or Th17-producing conditions, an effect was considered “common” using the criteria FDR < 0.05 in at least one comparison and (unadjusted) p-value < 0.05 in the other with a log fold-change in the same direction. Opposite effects satisfied the same significance criteria but with a log fold-change of opposing signs, and Th1/Th17-specific effects satisfied FDR < 0.05 in one group but p-value > 0.05 in the other.

### Analysis of in-vivo 10x-sequenced single-cell libraries

#### Alignment and Preprocessing

To quantify gene expression, samples were individually aligned to the mm10 mus musculus genome using CellRanger v2.0.2. In total, 14 samples were collected, however two samples of intraepithelial lymphocytes in the small intestine from batch 1 were discarded as the IL23R knockout sample did not pass quality checks (insufficient barcodes belonging to cells). The remaining 12 samples were used in downstream analysis.

#### Merging and Filtering

An initial analysis was performed on the concatenation of all samples to identify and remove dying or contaminating cell types. All 12 samples were modeled with scVI (Lopez et al., 2018), selecting genes expressed in at least 100 cells, using 10 latent components, and encoding each sample ID as a batch variable so as to remove tissue and genotype specific effects.

The resulting latent space was then clustered into 11 clusters using the Louvain algorithm as implemented in SCANPY (Wolf et al., 2018) with the following settings: n_neighbors = 10 and resolution = 0.5. Of these four clusters were marked for removal in all subsequent analysis (across all samples), in total consisting of 7.6% of all 32,763 cells.

One cluster, c8, was characterized by low recovered UMI counts, a low proportion of ribosomal coding RNA, a high proportion of mitochondrial RNA, and very few other distinguishing genes. These were removed as being likely low-quality measurements from damaged cells.

Another cluster, c9, was identified as a possible contaminant cell type due to marked lack of CD4 and CD52 expression. Similarly, cluster c10 lacked T cell markers and had high expression of hemoglobin-related genes.

Finally, cluster c6, despite exhibiting Th17-related genes (Ccr6, Il-23r, Rorc, and IL-22), was removed as the lack of CD3 expression, lack of TCR expression, and distinct lack of any IL-23r-eGFP detection (despite significant Il-23r detection), indicated that these were likely LTi ILC3 cells from the recipient mouse and therefore not part of the genotype-perturbed donor cells of interest in this experiment. After this stage of filtering, a total of 30,259 cells remained for downstream analysis.

#### Between Tissue Analyses

The combined UMAP plots of Figure 2 were generated by first modeling all cells with scVI (Lopez et al., 2018), and then running the UMAP (Becht et al., 2018) algorithm using SCANPY (Wolf et al., 2018) with n_neighbors=30. When running scVI, the two experimental batches were encoded as the batch variable and zero-inflation in the model was disabled. Additionally, genes were pre-filtered to retain A) genes expressed in at least 100 cells and B) genes with high fano factor: genes divided into 30 bins based on mean, fano factor computed for each gene - using the UMI-depth normalized counts - and genes retained with a fano factor greater than 2 MAD (median absolute deviations above median) per bin.

For the heatmap in Figure 3D, edgeR (Robinson et al., 2010) was run to compute average expression per tissue and to determine the genes with the highest tissue specificity, the top 500 of which were plotted. The model was fit in the form “Batch + Tissue” and the Tissue coefficient was tested with the likelihood ratio test. When computing within-tissue differential expression based on Genotype, edgeR was used to compute differential expression (between control and eGFP/eGFP cells) using the likelihood ratio test.

#### Within-Tissue Analysis: Lamina Propria Lymphocytes

Samples from the lamina propria (6 total - batch 1: control and knockout in colon and batch: 2 control and knockout in colon and small intestine) were grouped together and modeled with scVI (Lopez et al., 2018) (8 components, zero-inflation enabled, genes retained that were expressed in 10 cells or more and the Fano filter as described in the previous section). As broad genotype-specific differences in expression (e.g., heat shock proteins), prevented cells from clustering together, we regressed out individual sample differences as if they were batch (i.e., encoded sample ID as the batch variable) and instead assessed the resulting clusters in terms of proportional representation (control vs. knockout) and within-cluster differential expression (control vs knockout). UMAP and Louvain clustering were run on the resulting 8 components as implemented in SCANPY (Wolf et al., 2018) with the following parameters: n_neighbors=30 and (for Louvain) resolution=0.9.

Changes in cluster composition were adjusted by the number of cells recovered using the following relation:

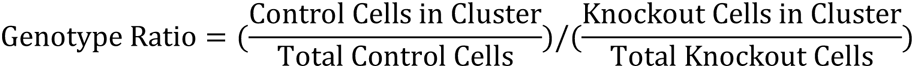

To identify marker genes per cluster, edgeR (Robinson et al., 2010) was used to compute log fold-change and significance when comparing cells in the cluster vs all other cells (1 vs. all differential expression). Similarly, edgeR was used to determine differential expression within cluster (control vs. knockout cells).

The Tr1 signature from Gruarin et al., 2019 (Gruarin et al., 2019) is composed of the 51 genes differentiated Tr1 cells from other CD4 cells in Figure 1A of that study. Per-cell signature scores were calculated using VISION (DeTomaso et al., 2019) and the expression matrix scaled to the median number of UMIs per cell.

#### Target Prioritization (Ranking)

For each gene, we computed an overall score with four components. Genes were then ranked by a weighted sum of these components. The components are defined as:

1. In-vivo, Cluster Specific - the maximum of:

- The log fold-change of genes significant (FDR < 0.1) in the cluster 2 vs. all comparison

- The log fold-change * 0.5 of genes significant (FDR < 0.1) in the cluster 9 vs. all comparison

- The log fold-change of genes significant (FDR < 0.1) in the within-cluster (cluster 2) control vs. knockout comparison

- The log fold-change * 0.5 of genes significant (FDR < 0.1) in the within-cluster (cluster 9) control vs. knockout comparison

2. In-vivo, Tissue Specific - the maximum of:

- The log fold-change for genes significant (FDR < 0.1) in the LPL vs. Spleen comparison

- The log fold-change for genes significant (FDR < 0.1) in the within-LPL control vs. knockout comparison

For both the above in-vivo categories, negative log fold-changes were truncated at 0 (to select specifically for genes upregulated in the cluster of interest or in control cells) and the overall score per gene was limited to 2.

3. In-vitro, Summary - the maximum of:

- The log fold-change for genes with significant (FDR < .1) genotype-independent differences in the in-vitro GFP+ vs. GFP-comparison

- The log fold-change of the genotype-dependent effect (GFP+ x Genotype Ctrl) in the in-vitro linear model

For the above, in-vitro category, the overall score per gene was limited to 3.

4. GWAS:

- We assigned a GWAS score of 1 for genes within GWAS loci for Ulcerative Colitis or Crohn’s Disease (as identified by Jostins et. al., 2012; Liu et. al., 2015; DeLange et. al., 2017 (de Lange et al., 2017; Jostins et al., 2012; Liu et al., 2015) and compiled by DeLange et al., 2017 (de Lange et al., 2017)). Alternate genes were discarded for loci in which a specific gene has been confidently implicated by fine-mapping, eQTL, or target sequencing studies (as compiled by DeLange et al. 2017 (de Lange et al., 2017)).

The final ranking was determined by combining all four categories using the following weightings:

Overall Score = (In-vivo, Cluster Specific) x 2 + (In-vivo, Tissue Specific) x 1 + (In-vitro, Summary) x 1 + (GWAS) x 2

## Supplemental Information titles and legends

**<u>Description of tables S1-S4 (in Excel format)</u>**

**Table S1**

Supplementary tables related to the Smart-Seq2, in-vitro T cell cytokine stimulation experiments. Includes cell-level meta-data (Stimulus/Genotype/GFP) along with the results of differential expression comparisons (GFP+ vs GFP-comparisons along with the results of the combined, two-factor model).

**Table S2**

Supplementary tables related to the 10x, single-cell sequencing on T cells derived from the in-vivo mouse experiments. Contains cell-level meta-data for all 32,764 sequenced cells along with the results of Tissue vs. Tissue differential expression comparisons (both between-tissue comparisons and within-tissue, genotype comparisons)

**Table S3**

Supplementary tables related to the 10x, single-cell sequencing on T cells derived from the in-vivo mouse experiments originating from the small intestine or colonic lamina propria. Contains the results of 1 vs all differential expression tests for each of the 12 clusters defined in the study. Additionally, contains the results of within-cluster, between-genotype differential expression comparisons.

**Table S4**

Supplementary table detailing the ranking procedure used to prioritize genes for follow-up study. Gathers, for each gene, the final score used for ranking in addition to the component statistics used in computing this score. Details on the formula used are provided in the supplementary methods.

## Notes

https://www.ncbi.nlm.nih.gov/geo/query/acc.cgi?acc=GSE164520

## References

Abadia-Molina, A.C., Ji, H., Faubion, W.A., Julien, A., Latchman, Y., Yagita, H., Sharpe, A., Bhan, A.K., and Terhorst, C. (2006). CD48 controls T-cell and antigen-presenting cell functions in experimental colitis. Gastroenterology 130, 424–434.

Acosta-Rodriguez, E.V., Napolitani, G., Lanzavecchia, A., and Sallusto, F. (2007). Interleukins 1beta and 6 but not transforming growth factor-beta are essential for the differentiation of interleukin 17-producing human T helper cells. Nature immunology 8, 942–949.

Ahern, P.P., Schiering, C., Buonocore, S., McGeachy, M.J., Cua, D.J., Maloy, K.J., and Powrie, F. (2010). Interleukin-23 drives intestinal inflammation through direct activity on T cells. Immunity 33, 279–288.

Alfen, J.S., Larghi, P., Facciotti, F., Gagliani, N., Bosotti, R., Paroni, M., Maglie, S., Gruarin, P., Vasco, C.M., Ranzani, V., et al. (2018). Intestinal IFN-gamma-producing type 1 regulatory T cells coexpress CCR5 and programmed cell death protein 1 and downregulate IL-10 in the inflamed guts of patients with inflammatory bowel disease. J Allergy Clin Immunol 142, 1537–1547 e1538.

Amarnath, S., Mangus, C.W., Wang, J.C., Wei, F., He, A., Kapoor, V., Foley, J.E., Massey, P.R., Felizardo, T.C., Riley, J.L., et al. (2011). The PDL1-PD1 axis converts human TH1 cells into regulatory T cells. Sci Transl Med 3, 111ra120.

Annacker, O., Coombes, J.L., Malmstrom, V., Uhlig, H.H., Bourne, T., Johansson-Lindbom, B., Agace, W.W., Parker, C.M., and Powrie, F. (2005). Essential role for CD103 in the T cell-mediated regulation of experimental colitis. J Exp Med 202, 1051–1061.

Anumanthan, A., Bensussan, A., Boumsell, L., Christ, A.D., Blumberg, R.S., Voss, S.D., Patel, A.T., Robertson, M.J., Nadler, L.M., and Freeman, G.J. (1998). Cloning of BY55, a novel Ig superfamily member expressed on NK cells, CTL, and intestinal intraepithelial lymphocytes. J Immunol 161, 2780–2790.

Atarashi, K., Suda, W., Luo, C., Kawaguchi, T., Motoo, I., Narushima, S., Kiguchi, Y., Yasuma, K., Watanabe, E., Tanoue, T., et al. (2017). Ectopic colonization of oral bacteria in the intestine drives TH1 cell induction and inflammation. Science 358, 359–365.

Awasthi, A., Riol-Blanco, L., Jager, A., Korn, T., Pot, C., Galileos, G., Bettelli, E., Kuchroo, V.K., and Oukka, M. (2009). Cutting edge: IL-23 receptor gfp reporter mice reveal distinct populations of IL-17-producing cells. J Immunol 182, 5904–5908.

Becattini, S., Latorre, D., Mele, F., Foglierini, M., De Gregorio, C., Cassotta, A., Fernandez, B., Kelderman, S., Schumacher, T.N., Corti, D., et al. (2015). T cell immunity. Functional heterogeneity of human memory CD4(+) T cell clones primed by pathogens or vaccines. Science 347, 400-406.

Becht, E., McInnes, L., Healy, J., Dutertre, C.A., Kwok, I.W.H., Ng, L.G., Ginhoux, F., and Newell, E.W. (2018). Dimensionality reduction for visualizing single-cell data using UMAP. Nat Biotechnol.

Bolger, A.M., Lohse, M., and Usadel, B. (2014). Trimmomatic: a flexible trimmer for Illumina sequence data. Bioinformatics 30, 2114–2120.

Bonecchi, R., Bianchi, G., Bordignon, P.P., D’Ambrosio, D., Lang, R., Borsatti, A., Sozzani, S., Allavena, P., Gray, P.A., Mantovani, A., and Sinigaglia, F. (1998). Differential expression of chemokine receptors and chemotactic responsiveness of type 1 T helper cells (Th1s) and Th2s. J Exp Med 187, 129–134.

Brass, A.L., Huang, I.C., Benita, Y., John, S.P., Krishnan, M.N., Feeley, E.M., Ryan, B.J., Weyer, J.L., van der Weyden, L., Fikrig, E., et al. (2009). The IFITM proteins mediate cellular resistance to influenza A H1N1 virus, West Nile virus, and dengue virus. Cell 139, 1243–1254.

Carlson, C.M., Endrizzi, B.T., Wu, J., Ding, X., Weinreich, M.A., Walsh, E.R., Wani, M.A., Lingrel, J.B., Hogquist, K.A., and Jameson, S.C. (2006). Kruppel-like factor 2 regulates thymocyte and T-cell migration. Nature 442, 299–302.

Cua, D.J., Sherlock, J., Chen, Y., Murphy, C.A., Joyce, B., Seymour, B., Lucian, L., To, W., Kwan, S., Churakova, T., et al. (2003). Interleukin-23 rather than interleukin-12 is the critical cytokine for autoimmune inflammation of the brain. Nature 421, 744–748.

Dardalhon, V., Schubart, A.S., Reddy, J., Meyers, J.H., Monney, L., Sabatos, C.A., Ahuja, R., Nguyen, K., Freeman, G.J., Greenfield, E.A., et al. (2005). CD226 is specifically expressed on the surface of Th1 cells and regulates their expansion and effector functions. J Immunol 175, 1558–1565.

de Lange, K.M., Moutsianas, L., Lee, J.C., Lamb, C.A., Luo, Y., Kennedy, N.A., Jostins, L., Rice, D.L., Gutierrez-Achury, J., Ji, S.G., et al. (2017). Genome-wide association study implicates immune activation of multiple integrin genes in inflammatory bowel disease. Nat Genet 49, 256–261.

DeTomaso, D., Jones, M.G., Subramaniam, M., Ashuach, T., Ye, C.J., and Yosef, N. (2019). Functional interpretation of single cell similarity maps. Nat Commun 10, 4376.

DiSpirito, J.R., Zemmour, D., Ramanan, D., Cho, J., Zilionis, R., Klein, A.M., Benoist, C., and Mathis, D. (2018). Molecular diversification of regulatory T cells in nonlymphoid tissues. Sci Immunol 3, eaat5861

Duerr, R.H., Taylor, K.D., Brant, S.R., Rioux, J.D., Silverberg, M.S., Daly, M.J., Steinhart, A.H., Abraham, C., Regueiro, M., Griffiths, A., et al. (2006). A genome-wide association study identifies IL23R as an inflammatory bowel disease gene. Science 314, 1461–1463.

Finak, G., McDavid, A., Yajima, M., Deng, J., Gersuk, V., Shalek, A.K., Slichter, C.K., Miller, H.W., McElrath, M.J., Prlic, M., et al. (2015). MAST: a flexible statistical framework for assessing transcriptional changes and characterizing heterogeneity in single-cell RNA sequencing data. Genome Biol 16, 278.

Forster, R., Davalos-Misslitz, A.C., and Rot, A. (2008). CCR7 and its ligands: balancing immunity and tolerance. Nat Rev Immunol 8, 362–371.

Fuchs, A., Vermi, W., Lee, J.S., Lonardi, S., Gilfillan, S., Newberry, R.D., Cella, M., and Colonna, M. (2013). Intraepithelial type 1 innate lymphoid cells are a unique subset of IL-12- and IL-15-responsive IFN-gamma-producing cells. Immunity 38, 769–781.

Gagliani, N., Amezcua Vesely, M.C., Iseppon, A., Brockmann, L., Xu, H., Palm, N.W., de Zoete, M.R., Licona-Limon, P., Paiva, R.S., Ching, T., et al. (2015). Th17 cells transdifferentiate into regulatory T cells during resolution of inflammation. Nature 523, 221–225.

Gaublomme, J.T., Yosef, N., Lee, Y., Gertner, R.S., Yang, L.V., Wu, C., Pandolfi, P.P., Mak, T., Satija, R., Shalek, A.K., et al. (2015). Single-Cell Genomics Unveils Critical Regulators of Th17 Cell Pathogenicity. Cell 163, 1400–1412.

Ghoreschi, K., Laurence, A., Yang, X.P., Tato, C.M., McGeachy, M.J., Konkel, J.E., Ramos, H.L., Wei, L., Davidson, T.S., Bouladoux, N., et al. (2010). Generation of pathogenic T(H)17 cells in the absence of TGF-beta signalling. Nature 467, 967–971.

Gilfillan, S., Chan, C.J., Cella, M., Haynes, N.M., Rapaport, A.S., Boles, K.S., Andrews, D.M., Smyth, M.J., and Colonna, M. (2008). DNAM-1 promotes activation of cytotoxic lymphocytes by nonprofessional antigen-presenting cells and tumors. J Exp Med 205, 2965–2973.

Grenningloh, R., Kang, B.Y., and Ho, I.C. (2005). Ets-1, a functional cofactor of T-bet, is essential for Th1 inflammatory responses. J Exp Med 201, 615–626.

Gruarin, P., Maglie, S., De Simone, M., Haringer, B., Vasco, C., Ranzani, V., Bosotti, R., Noddings, J.S., Larghi, P., Facciotti, F., et al. (2019). Eomesodermin controls a unique differentiation program in human IL-10 and IFN-gamma coproducing regulatory T cells. Eur J Immunol 49, 96–111.

Harbour, S.N., Maynard, C.L., Zindl, C.L., Schoeb, T.R., and Weaver, C.T. (2015). Th17 cells give rise to Th1 cells that are required for the pathogenesis of colitis. Proc Natl Acad Sci U S A 112, 7061–7066.

Hirota, K., Duarte, J.H., Veldhoen, M., Hornsby, E., Li, Y., Cua, D.J., Ahlfors, H., Wilhelm, C., Tolaini, M., Menzel, U., et al. (2011). Fate mapping of IL-17-producing T cells in inflammatory responses. Nat Immunol 12, 255–263.

Hsu, C.Y., Yeh, L.T., Fu, S.H., Chien, M.W., Liu, Y.W., Miaw, S.C., Chang, D.M., and Sytwu, H.K. (2018). SUMO-defective c-Maf preferentially transactivates Il21 to exacerbate autoimmune diabetes. J Clin Invest 128, 3779–3793.

Huang, H., Fang, M., Jostins, L., Umicevic Mirkov, M., Boucher, G., Anderson, C.A., Andersen, V., Cleynen, I., Cortes, A., Crins, F., et al. (2017). Fine-mapping inflammatory bowel disease loci to single-variant resolution. Nature 547, 173–178.

Hueber, W., Sands, B.E., Lewitzky, S., Vandemeulebroecke, M., Reinisch, W., Higgins, P.D., Wehkamp, J., Feagan, B.G., Yao, M.D., Karczewski, M., et al. (2012). Secukinumab, a human anti-IL-17A monoclonal antibody, for moderate to severe Crohn’s disease: unexpected results of a randomised, double-blind placebo-controlled trial. Gut 61, 1693–1700.

Ito, R., Shin-Ya, M., Kishida, T., Urano, A., Takada, R., Sakagami, J., Imanishi, J., Kita, M., Ueda, Y., Iwakura, Y., et al. (2006). Interferon-gamma is causatively involved in experimental inflammatory bowel disease in mice. Clin Exp Immunol 146, 330–338.

Iwamoto, T., Suto, A., Tanaka, S., Takatori, H., Suzuki, K., Iwamoto, I., and Nakajima, H. (2014). Interleukin-21-producing c-Maf-expressing CD4+ T cells induce effector CD8+ T cells and enhance autoimmune inflammation in scurfy mice. Arthritis Rheumatol 66, 2079–2090.

Izcue, A., Hue, S., Buonocore, S., Arancibia-Carcamo, C.V., Ahern, P.P., Iwakura, Y., Maloy, K.J., and Powrie, F. (2008). Interleukin-23 restrains regulatory T cell activity to drive T cell-dependent colitis. Immunity 28, 559–570.

Jackson, R., Kroehling, L., Khitun, A., Bailis, W., Jarret, A., York, A.G., Khan, O.M., Brewer, J.R., Skadow, M.H., Duizer, C., et al. (2018). The translation of non-canonical open reading frames controls mucosal immunity. Nature 564, 434–438.

Johansson-Lindbom, B., and Agace, W.W. (2007). Generation of gut-homing T cells and their localization to the small intestinal mucosa. Immunol Rev 215, 226–242.

Jostins, L., Ripke, S., Weersma, R.K., Duerr, R.H., McGovern, D.P., Hui, K.Y., Lee, J.C., Schumm, L.P., Sharma, Y., Anderson, C.A., et al. (2012). Host-microbe interactions have shaped the genetic architecture of inflammatory bowel disease. Nature 491, 119–124.

Kakiuchi, N., Yoshida, K., Uchino, M., Kihara, T., Akaki, K., Inoue, Y., Kawada, K., Nagayama, S., Yokoyama, A., Yamamoto, S., et al. (2020). Frequent mutations that converge on the NFKBIZ pathway in ulcerative colitis. Nature 577, 260–265.

Khor, B., Gardet, A., and Xavier, R.J. (2011). Genetics and pathogenesis of inflammatory bowel disease. Nature 474, 307–317.

Korn, T., Bettelli, E., Gao, W., Awasthi, A., Jager, A., Strom, T.B., Oukka, M., and Kuchroo, V.K. (2007). IL-21 initiates an alternative pathway to induce proinflammatory T(H)17 cells. Nature 448, 484–487.

Kowalczyk, M.S., Tirosh, I., Heckl, D., Rao, T.N., Dixit, A., Haas, B.J., Schneider, R.K., Wagers, A.J., Ebert, B.L., and Regev, A. (2015). Single-cell RNA-seq reveals changes in cell cycle and differentiation programs upon aging of hematopoietic stem cells. Genome Res 25, 1860–1872.

Kurdi, A.T., Bassil, R., Olah, M., Wu, C., Xiao, S., Taga, M., Frangieh, M., Buttrick, T., Orent, W., Bradshaw, E.M., et al. (2016). Tiam1/Rac1 complex controls Il17a transcription and autoimmunity. Nat Commun 7, 13048.

Lamb, C.A., Mansfield, J.C., Tew, G.W., Gibbons, D., Long, A.K., Irving, P., Diehl, L., Eastham-Anderson, J., Price, M.B., O’Boyle, G., et al. (2017). alphaEbeta7 Integrin Identifies Subsets of Pro-Inflammatory Colonic CD4+ T Lymphocytes in Ulcerative Colitis. J Crohns Colitis 11, 610–620.

Langmead, B., and Salzberg, S.L. (2012). Fast gapped-read alignment with Bowtie 2. Nat Methods 9, 357–359.

Laudato, S., Patil, N., Abba, M.L., Leupold, J.H., Benner, A., Gaiser, T., Marx, A., and Allgayer, H. (2017). P53-induced miR-30e-5p inhibits colorectal cancer invasion and metastasis by targeting ITGA6 and ITGB1. Int J Cancer 141, 1879–1890.

Lee, Y., Awasthi, A., Yosef, N., Quintana, F.J., Xiao, S., Peters, A., Wu, C., Kleinewietfeld, M., Kunder, S., Hafler, D.A., et al. (2012a). Induction and molecular signature of pathogenic TH17 cells. Nat Immunol 13, 991–999.

Lee, Y., Awasthi, A., Yosef, N., Quintana, F.J., Xiao, S., Peters, A., Wu, C., Kleinewietfeld, M., Kunder, S., Hafler, D.A., et al. (2012b). Induction and molecular signature of pathogenic TH17 cells. Nature immunology 13, 991–999.

Li, B., and Dewey, C.N. (2011). RSEM: accurate transcript quantification from RNA-Seq data with or without a reference genome. BMC Bioinformatics 12, 323.

Liu, J.Z., van Sommeren, S., Huang, H., Ng, S.C., Alberts, R., Takahashi, A., Ripke, S., Lee, J.C., Jostins, L., Shah, T., et al. (2015). Association analyses identify 38 susceptibility loci for inflammatory bowel disease and highlight shared genetic risk across populations. Nat Genet 47, 979–986.

Lopez, R., Regier, J., Cole, M.B., Jordan, M.I., and Yosef, N. (2018). Deep generative modeling for single-cell transcriptomics. Nat Methods 15, 1053–1058.

Mailand, N., and Diffley, J.F. (2005). CDKs promote DNA replication origin licensing in human cells by protecting Cdc6 from APC/C-dependent proteolysis. Cell 122, 915–926.

Masopust, D., and Soerens, A.G. (2019). Tissue-Resident T Cells and Other Resident Leukocytes. Annu Rev Immunol 37, 521–546.

McGeachy, M.J., Chen, Y., Tato, C.M., Laurence, A., Joyce-Shaikh, B., Blumenschein, W.M., McClanahan, T.K., O’Shea, J.J., and Cua, D.J. (2009). The interleukin 23 receptor is essential for the terminal differentiation of interleukin 17-producing effector T helper cells in vivo. Nature immunology 10, 314–324.

Mosmann, T.R., Cherwinski, H., Bond, M.W., Giedlin, M.A., and Coffman, R.L. (1986). Two types of murine helper T cell clone. I. Definition according to profiles of lymphokine activities and secreted proteins. J Immunol 136, 2348–2357.

Murphy, C.A., Langrish, C.L., Chen, Y., Blumenschein, W., McClanahan, T., Kastelein, R.A., Sedgwick, J.D., and Cua, D.J. (2003). Divergent pro- and antiinflammatory roles for IL-23 and IL-12 in joint autoimmune inflammation. J Exp Med 198, 1951–1957.

Nurieva, R., Yang, X.O., Martinez, G., Zhang, Y., Panopoulos, A.D., Ma, L., Schluns, K., Tian, Q., Watowich, S.S., Jetten, A.M., and Dong, C. (2007). Essential autocrine regulation by IL-21 in the generation of inflammatory T cells. Nature 448, 480–483.

Patel, D.D., and Kuchroo, V.K. (2015). Th17 Cell Pathway in Human Immunity: Lessons from Genetics and Therapeutic Interventions. Immunity 43, 1040–1051.

Picelli, S., Bjorklund, A.K., Faridani, O.R., Sagasser, S., Winberg, G., and Sandberg, R. (2013). Smart-seq2 for sensitive full-length transcriptome profiling in single cells. Nat Methods 10, 1096–1098.

Powrie, F., Leach, M.W., Mauze, S., Menon, S., Caddle, L.B., and Coffman, R.L. (1994). Inhibition of Th1 responses prevents inflammatory bowel disease in scid mice reconstituted with CD45RBhi CD4+ T cells. Immunity 1, 553–562.

Robinson, M.D., McCarthy, D.J., and Smyth, G.K. (2010). edgeR: a Bioconductor package for differential expression analysis of digital gene expression data. Bioinformatics 26, 139–140.

Roman, J., Planell, N., Lozano, J.J., Aceituno, M., Esteller, M., Pontes, C., Balsa, D., Merlos, M., Panes, J., and Salas, A. (2013). Evaluation of responsive gene expression as a sensitive and specific biomarker in patients with ulcerative colitis. Inflamm Bowel Dis 19, 221–229.

Sandborn, W.J., Gasink, C., Gao, L.L., Blank, M.A., Johanns, J., Guzzo, C., Sands, B.E., Hanauer, S.B., Targan, S., Rutgeerts, P., et al. (2012). Ustekinumab induction and maintenance therapy in refractory Crohn’s disease. N Engl J Med 367, 1519–1528.

Sands, B.E., Peyrin-Biroulet, L., Loftus, E.V., Jr., Danese, S., Colombel, J.F., Toruner, M., Jonaitis, L., Abhyankar, B., Chen, J., Rogers, R., et al. (2019a). Vedolizumab versus Adalimumab for Moderate-to-Severe Ulcerative Colitis. N Engl J Med 381, 1215–1226.

Sands, B.E., Sandborn, W.J., Panaccione, R., O’Brien, C.D., Zhang, H., Johanns, J., Adedokun, O.J., Li, K., Peyrin-Biroulet, L., Van Assche, G., et al. (2019b). Ustekinumab as Induction and Maintenance Therapy for Ulcerative Colitis. N Engl J Med 381, 1201–1214.

Sato, K., Miyoshi, F., Yokota, K., Araki, Y., Asanuma, Y., Akiyama, Y., Yoh, K., Takahashi, S., Aburatani, H., and Mimura, T. (2011). Marked induction of c-Maf protein during Th17 cell differentiation and its implication in memory Th cell development. J Biol Chem 286, 14963–14971.

Schiering, C., Krausgruber, T., Chomka, A., Frohlich, A., Adelmann, K., Wohlfert, E.A., Pott, J., Griseri, T., Bollrath, J., Hegazy, A.N., et al. (2014). The alarmin IL-33 promotes regulatory T-cell function in the intestine. Nature 513, 564–568.

Schneider, M., Schumacher, V., Lischke, T., Lucke, K., Meyer-Schwesinger, C., Velden, J., Koch-Nolte, F., and Mittrucker, H.W. (2015). CD38 is expressed on inflammatory cells of the intestine and promotes intestinal inflammation. PLoS One 10, e0126007.

Singh, U.P., Singh, S., Taub, D.D., and Lillard, J.W., Jr. (2003). Inhibition of IFN-gamma-inducible protein-10 abrogates colitis in IL-10-/-mice. J Immunol 171, 1401–1406.

Szabo, S.J., Kim, S.T., Costa, G.L., Zhang, X., Fathman, C.G., and Glimcher, L.H. (2000). A novel transcription factor, T-bet, directs Th1 lineage commitment. Cell 100, 655-669.

Takahashi, S., Andreoletti, G., Chen, R., Munehira, Y., Batra, A., Afzal, N.A., Beattie, R.M., Bernstein, J.A., Ennis, S., and Snyder, M. (2017). De novo and rare mutations in the HSPA1L heat shock gene associated with inflammatory bowel disease. Genome Med 9, 8.

Tan, C.L., Peluso, M.J., Drijvers, J.M., Mera, C.M., Grande, S.M., Brown, K.E., Godec, J., Freeman, G.J., and Sharpe, A.H. (2018). CD160 Stimulates CD8(+) T Cell Responses and Is Required for Optimal Protective Immunity to Listeria monocytogenes. Immunohorizons 2, 238–250.

Teng, M.W., Bowman, E.P., McElwee, J.J., Smyth, M.J., Casanova, J.L., Cooper, A.M., and Cua, D.J. (2015). IL-12 and IL-23 cytokines: from discovery to targeted therapies for immune-mediated inflammatory diseases. Nat Med 21, 719–729.

Tu, T.C., Brown, N.K., Kim, T.J., Wroblewska, J., Yang, X., Guo, X., Lee, S.H., Kumar, V., Lee, K.M., and Fu, Y.X. (2015). CD160 is essential for NK-mediated IFN-gamma production. J Exp Med 212, 415–429.

Uniken Venema, W.T., Voskuil, M.D., Vila, A.V., van der Vries, G., Jansen, B.H., Jabri, B., Faber, K.N., Dijkstra, G., Xavier, R.J., Wijmenga, C., et al. (2019). Single-Cell RNA Sequencing of Blood and Ileal T Cells From Patients With Crohn’s Disease Reveals Tissue-Specific Characteristics and Drug Targets. Gastroenterology 156, 812–815 e822.

van der Maaten, L. (2014). Accelerating t-SNE using Tree-Based Algorithms. Journal of Machine Learning Research 15, 3221–3245.

van Eden, W., van der Zee, R., and Prakken, B. (2005). Heat-shock proteins induce T-cell regulation of chronic inflammation. Nat Rev Immunol 5, 318–330.

Velasquez, S., Malik, S., Lutz, S.E., Scemes, E., and Eugenin, E.A. (2016). Pannexin1 Channels Are Required for Chemokine-Mediated Migration of CD4+ T Lymphocytes: Role in Inflammation and Experimental Autoimmune Encephalomyelitis. J Immunol 196, 4338–4347.

Wang, X., Sumida, H., and Cyster, J.G. (2014). GPR18 is required for a normal CD8alphaalpha intestinal intraepithelial lymphocyte compartment. J Exp Med 211, 2351–2359.

Weder, B., Mozaffari, M., Biedermann, L., Mamie, C., Moncsek, A., Wang, L., Clarke, S.H., Rogler, G., McRae, B.L., Graff, C.L., et al. (2018). BCL-2 levels do not predict azathioprine treatment response in inflammatory bowel disease, but inhibition induces lymphocyte apoptosis and ameliorates colitis in mice. Clin Exp Immunol 193, 346–360.

Wells, M.L., Perera, L., and Blackshear, P.J. (2017). An Ancient Family of RNA-Binding Proteins: Still Important! Trends Biochem Sci 42, 285–296.

Wolf, F.A., Angerer, P., and Theis, F.J. (2018). SCANPY: large-scale single-cell gene expression data analysis. Genome Biol 19, 15.

Wu, C., Chen, Z., Xiao, S., Thalhamer, T., Madi, A., Han, T., and Kuchroo, V. (2018). SGK1 Governs the Reciprocal Development of Th17 and Regulatory T Cells. Cell Rep 22, 653–665.

Yu, H. (2007). Cdc20: a WD40 activator for a cell cycle degradation machine. Mol Cell 27, 3–16.

Zhang, P., Lee, J.S., Gartlan, K.H., Schuster, I.S., Comerford, I., Varelias, A., Ullah, M.A., Vuckovic, S., Koyama, M., Kuns, R.D., et al. (2017). Eomesodermin promotes the development of type 1 regulatory T (TR1) cells. Sci Immunol 2, eaah7152.

Zhang, Q., and Vignali, D.A. (2016). Co-stimulatory and Co-inhibitory Pathways in Autoimmunity. Immunity 44, 1034–1051.

Zhou, L., Ivanov, II, Spolski, R., Min, R., Shenderov, K., Egawa, T., Levy, D.E., Leonard, W.J., and Littman, D.R. (2007). IL-6 programs T(H)-17 cell differentiation by promoting sequential engagement of the IL-21 and IL-23 pathways. Nature immunology 8, 967–974.

